# Insights into Genome Ejection by a Therapeutic phiKMV-like Bacteriophage

**DOI:** 10.64898/2026.05.18.726046

**Authors:** Junwei Li, Nathan F. Bellis, Chun-Feng David Hou, Steven Branston, Zsuzsanna Kovach, Renae Geier, Angela Soriaga, Lucy Sim, Pierre Kyme, Deborah Birx, Sebastien Lemire, Gino Cingolani

**Author notes:** Corresponding Author: Gino Cingolani, Ph.D., Department of Biochemistry and Molecular Genetics, Univ of Alabama at Birmingham, 1825 University Blvd, Birmingham, AL 35294, USA. These authors contributed equally to this work (Junwei Li, Nathan F. Bellis).

## Abstract

Ar-KM is a phiKMV-like therapeutic bacteriophage used in clinical candidate phage therapy cocktails to treat lung infections caused by Pseudomonas aeruginosa. Here, we present an integrative structural atlas of Ar-KM proteins using cryo-EM, proteomics, and bioinformatics. From a single purified Ar-KM preparation, we identified three distinct populations: mature DNA-filled virions, open-nozzle particles with ejection proteins extending from the tail, and closed-nozzle empty particles. Near-atomic-resolution reconstructions of all three states enabled us to build atomic models for eleven structural proteins. The mature virion revealed the pre-ejection conformation of three ejection proteins, gp41, gp42, and gp43, homologous to coliphage T7’s gp14, gp15, and gp16, respectively. Unlike T7, peptidoglycan hydrolase activity associated with the ejectosome resides in the gp15-like periplasmic tunnel protein gp42, whereas in T7 the lysozyme-like domain is located at the N-terminus of gp16, underscoring the structural plasticity and evolutionary mosaicity of ejection proteins. We further identified a short α-helical factor, gp34, present in eight copies at the mismatched interface between the portal barrel and gp41. Gp34 forms a cage within the nozzle, acting as a molecular wedge that stabilizes the open conformation and permits gp41 to assemble into a hexameric channel during ejection. Evolutionarily, gp34 appears to be an ortholog of the essential gene gp7.3 in phage T7 and is conserved across sequenced phiKMV-like phages. We propose that this protein functions as an ejection protein assembly factor, stabilizing the open nozzle during infection and allowing the coordinated exit of ejection proteins and their assembly into a DNA-ejectosome.

**Highlights:**

- Structural polymorphism of phage Ar-KM defines open and closed nozzle states
- Ejection proteins gp43, gp42, and gp41 assemble in a 4:8:8 stoichiometry
- Lysozyme domain shows modular positioning within the ejectosome
- Portal protein acts as the primary barrier to genome leakage
- Wedge protein gp34 (T7 gp7.3 ortholog) bridges portal and gp41

## INTRODUCTION

*Pseudomonas aeruginosa* is an opportunistic pathogen that causes severe acute and chronic infections, particularly in individuals with cystic fibrosis and non-cystic fibrosis bronchiectasis. Rising antimicrobial resistance has intensified interest in bacteriophage therapy as an alternative or adjunctive treatment strategy [1, 2]. Among the diverse bacterial viruses that infect *P. aeruginosa*, phages of the *Autographiviridae* family have emerged as promising candidates due to their typically lytic lifestyle, rapid infection cycle, and compact genomes [3]. PhiKMV, a podophage and the founding member of the *Phikmvvirus* genus, is a lytic virus that infects *P. aeruginosa*. Initially discovered in 2013 [4] and distantly related to phage T7, phiKMV possesses its own RNA polymerase to express its late genes. PhiKMV features a linear, GC-rich (∼62.3%) double-stranded DNA genome spanning approximately 42kb, flanked by 414-base-pair direct terminal repeats and containing 48 open reading frames (ORFs), all transcribed from the same strand. Although its DNA shows little homology to known phages, 11 of its proteins share sequence similarity with T7-like phage proteins [4]. Among the 48 ORFs, 18 have been functionally annotated, including RNA polymerase, proteins involved in DNA replication, structural components, maturation factors, and lysis-related proteins. However, mass spectrometry (MS) analysis of purified phiKMV virions revealed more than 20 structural proteins [5], suggesting additional ORFs may be present in the capsid and ejected during infection.

The mechanisms by which *Phikmvviruses* infect *P. aeruginosa* and deliver their genomes into the bacterium are poorly understood. Phage phiKMV has been shown to require the host’s type IV pili (T4P) for successful infection: loss of the major pilin gene (pilA) renders the host resistant to phiKMV, whereas complementation restores both twitching motility and phage sensitivity, demonstrating a direct link between T4P expression and phiKMV adsorption [6–10]. Similarly, Luz19, a virulent *Phikmvvirus* of *P. aeruginosa*, has been characterized as T4P-dependent, with phage-resistant bacterial variants often carrying mutations in T4P-associated genes [11]. This receptor usage appears to be shared across many members of the broader phiKMV/Luz19-like phage group, although the T4P is unlikely to be the only receptor for phiKMV-like phages. It was hypothesized [7] that the pilus serves as a fishing hook for the phages, bringing them into close proximity to the cell surface upon retraction, thereby facilitating the interaction with a secondary receptor, likely deep within the core LPS, to trigger irreversible adsorption and DNA ejection. Despite the dependence on T4P as the primary receptor, evidence for a defined secondary receptor is lacking for both phiKMV and Luz19 [4]. In studies with phiKMV-related phages, screening of outer membrane (OM) mutants did not reveal a consistent additional receptor beyond T4P, and differences in host range among clinical isolates were attributed to variation in intracellular development rather than to alternate adsorption mechanisms [12]. A limitation of these studies is that LPS is often essential, and mutations that affect its structure are likely deleterious. Similarly, for Luz19, no alternative receptor has been described; phage-resistant variants lose susceptibility primarily through loss or alteration of T4P, without compensatory changes to lipopolysaccharide (LPS) or other OM components [13]. Therefore, in *P. aeruginosa* the current evidence supports a model in which phiKMV and Luz19 infect via a single, T4P-dependent adsorption step, with no confirmed secondary receptor.

Unlike T4P-specific phages such as phiKMV, phage T7 uses the host LPS as its primary (and likely only) receptor for *attachment to Escherichia coli (E. coli)*. Gene product gp17 is the tail fiber protein [14] that mediates initial, reversible adsorption by binding the LPS core oligosaccharide on the *E. coli* surface. This low-affinity interaction tethers the virion and promotes surface diffusion, facilitating subsequent irreversible attachment of the nozzle [15], which triggers tail rearrangements and causes the ejection of internal virion proteins, gp14, gp15, and gp16, into the bacterium’s cell envelope. Here, the three ejection proteins form a trans-envelope channel that enables efficient genome delivery into the host [16]. Before ejection, gp14, gp15, and gp16 form coaxially stacked rings onto the portal protein [17–19]. After infection, they assemble into a transmembrane channel within the bacterial cell envelope [20], or DNA-ejectosome, where gp14 forms an outer membrane complex (OMC), gp15 folds into a periplasmic tunnel (PT) connecting the two membranes [21, 22], and gp16 assembles as an inner membrane complex (IMC) that recruits the host’s RNA polymerase (RNAP) to pull the viral genome into the cell [23]. However, not all podophages have ejection proteins organized within the virion prior to ejection [20, 24]. For instance, in Lederberg viruses such as P22 [25] and Sf6 [26], ejection proteins are not visible by cryo-EM asymmetric analysis [27]. Indirect evidence suggests these proteins occupy space within the virion, perhaps loosely associated with the portal barrel [27]. In other phages, such as the *Litunavirus* DEV [28] and the *Bruynoghevirus* Pa223 [29], the gp15-like ejection protein is located around the perimeter of the portal protein before ejection and assembles into a nonameric PT after ejection. In contrast, *Litunavirus* phages such as DEV and N4 encode a large (∼3,500-residue) gp16-like ejection protein, also known as the virion-associated RNA polymerase (vRNAP) [30, 31].

Multiple studies have assessed phiKMV and related phages in therapeutic contexts [32]. Bioinformatics evidence suggests that phiKMV-like phages do not encode any known bacterial toxins or virulence factors that could pose a risk to humans. *In vitro*, phiKMV exhibits potent lytic activity against diverse *P. aeruginosa* isolates and produces halos on bacterial lawns, suggesting depolymerase or oligosaccharide-degrading activity, although it cannot degrade alginate produced by mucoid cystic fibrosis strains due to the lack of alginate-degrading enzymes [33]. In this paper, we study *Pseudomonas* phage Ar-KM (Armata phiKMV-like), a phiKMV-like phage isolated by Armata Pharmaceuticals and used for its bactericidal activity. The Ar-KM genome is 87% identical to Luz19 and 89% identical to phiKMV (Table S1), and it contains conserved morphogenesis-related genes. Deletion of the major pilin gene (pilA) along with many other pilus-related mutations (*pilE*, *pilB*, *pilQ*, *pilD*, *pilM*, *pilW*) make the host resistant to Ar-KM, indicating that the phage attaches via the T4P and requires its ability to extend and retract (*personal communication*), as expected given its high similarity to phiKMV and Luz19. Ar-KM was recently included in a therapeutic cocktail of five phages used by Armata Pharmaceuticals to treat *P. aeruginosa* infections in CF patients and those with non-CF bronchiectasis, as part of clinical trials SWARM-P.a. (NCT04596319) and Tailwind (NCT05616221), respectively.

In this study, we present a comprehensive structural atlas of the therapeutic phage Ar-KM. Our findings provide mechanistic insight into how Ar-KM gates its genome inside the capsid, attaches to the T4P, and delivers ejection proteins into the bacterium. This research expands our understanding of the architecture and dynamics of the ejection machinery before and during infection in a T4P-specific phage.

## RESULTS

### Cryo-EM analysis reveals three populations of phage Ar-KM

We vitrified *Pseudomonas* phage Ar-KM and imaged cryo-grids on a Titan Krios transmission electron microscope at 300 kV (Fig. 1A, Table 1). Initial 2D classification revealed two populations comprising ∼17,000 DNA-filled virions (FV) and ∼10,000 empty particles (EP) (Fig. S1). Both capsids were fully angular with a small tail apparatus characteristic of podoviruses. However, empty particles lacked bulk DNA inside the capsid. We computed icosahedral reconstructions for both species and applied 6-fold rotational symmetry (C6) to the tail region. Strikingly, FV had a large density above the portal protein, similar to that observed in phage T7 [17], which we improved by applying 4-fold rotational symmetry (C4) (Fig. 1B, Fig. S2). Additionally, 3D classification of EP along the tail revealed two tail apparatus conformations, which we separated and subjected to local refinement, imposing C6 symmetry (Fig. S1). Notably, one population (6,711 particles) had an open nozzle, with additional density lining the lumen (Fig. 1C), hereafter referred to as the ejecting particle (EJP). The smaller population (3,004 particles) consisted of empty particles (EP) with a closed nozzle (Fig. 1D). Thus, from a single purified preparation, we reconstructed three distinct structural states of phage Ar-KM, FV, EJP, and EP, at near-atomic resolution (2.8–3.2 Å; Table 1, Fig. S3). Well-defined side-chain density (Fig. S4) allowed us to annotate and *de novo* build eleven Ar-KM ORFs, which collectively were real-space refined to a map-to-model correlation coefficient (CC) ranging from 0.81 to 0.90 (Fig. 2, Table 1). We also identified and generated AlphaFold 3 models for three additional tail factors, gp45, gp46, and gp47, for which the experimental electron density is weak. LC-MS/MS analysis of virions used for cryo-EM confirmed the presence of all proteins identified in the cryo-EM density. It also revealed gp36, a putative scaffolding protein that was not readily visible in the reconstructions (Table 2).

**Figure 1.**
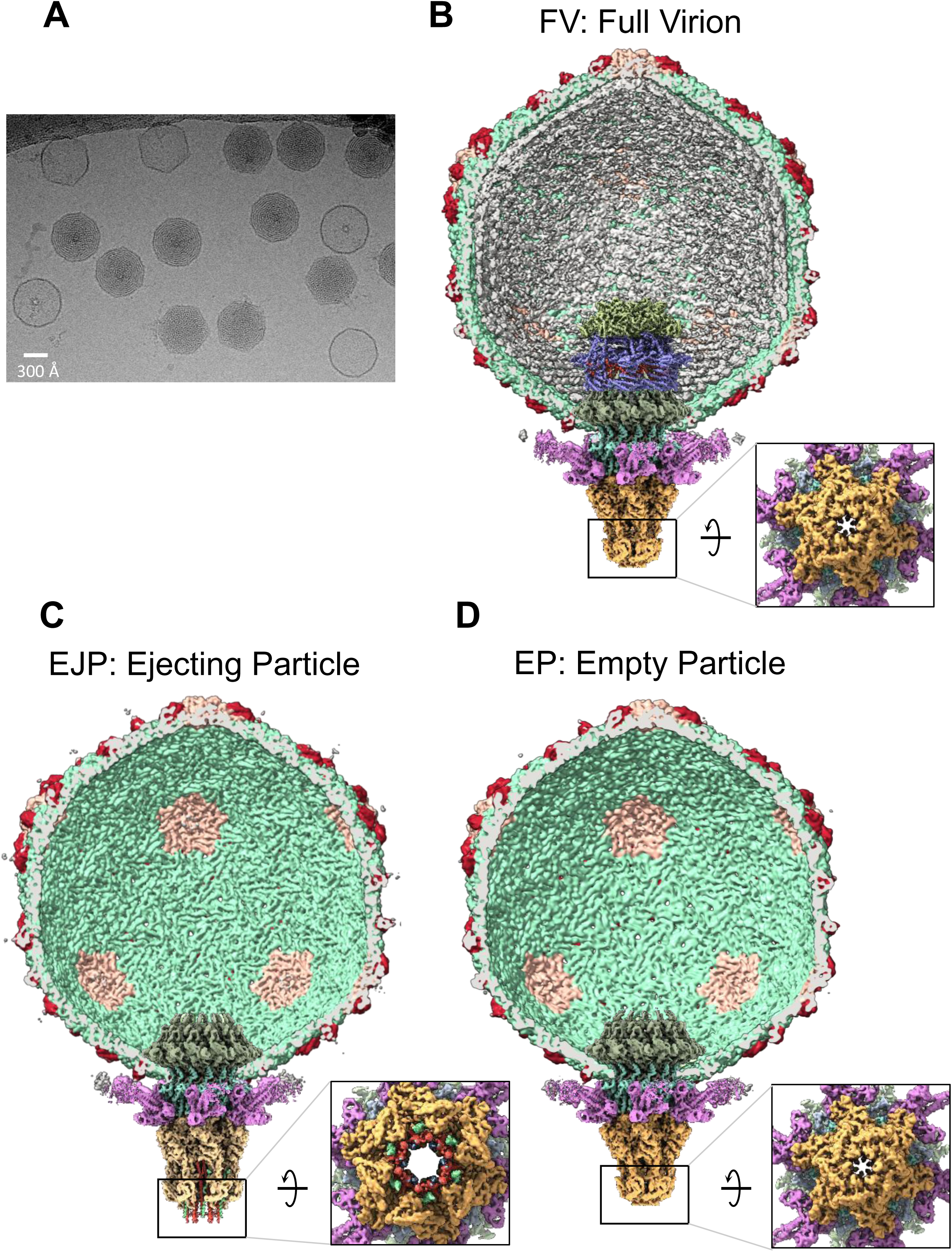
Cryo-EM reconstructions of *Pseudomonas* phage Ar-KM. (A) A representative cryo-micrograph of vitrified Ar-KM. (B-D) Composite reconstructions of the three Ar-KM populations derived from the same dataset. In (B), a cutaway view of the full virion (FV) of Ar-KM shows ordered ejection proteins stacked onto the portal protein and a closed nozzle. (C) Ar-KM ejecting particles (EJP) with an empty capsid and ejection proteins extending from the open tail nozzle. (D) Ar-KM empty particles (EP) that have released both DNA and ejection proteins, with a closed tail nozzle.

**Figure 2.**
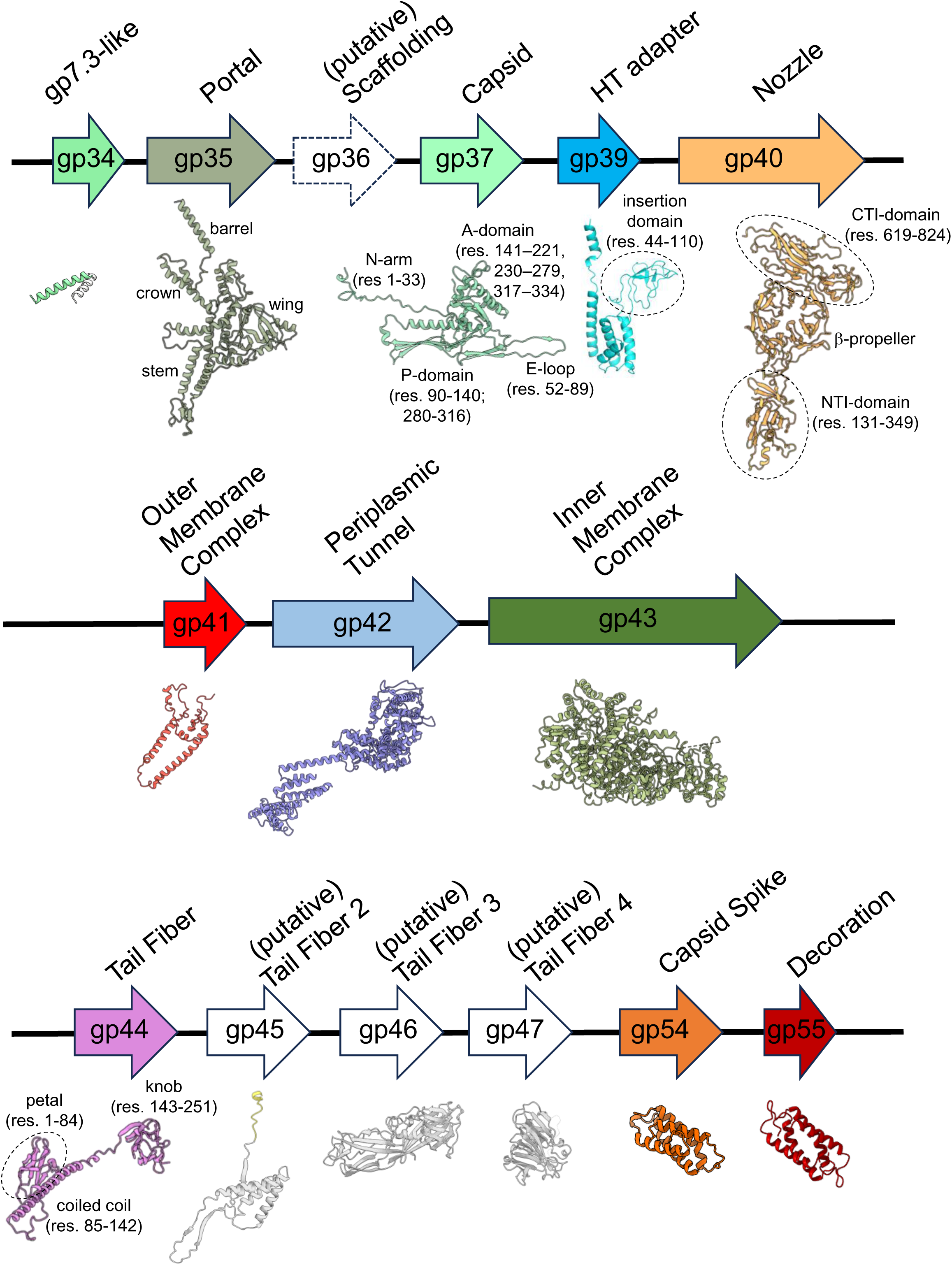
Organization of Ar-KM structural genes. Schematic diagram of the gene products of phage Ar-KM that encode structural proteins identified in this study. Ribbon diagrams for each structural protein, built de novo in this study, are shown below the gene product and are color-coded consistently with the other figures presented in this paper. Putative gene products not directly visualized by cryo-EM but detected by MS (Table 2) were predicted with AlphaFold and are shown in white, except for gp36 (putative scaffolding protein), for which the AlphaFold models have low confidence.

**Table 1.**
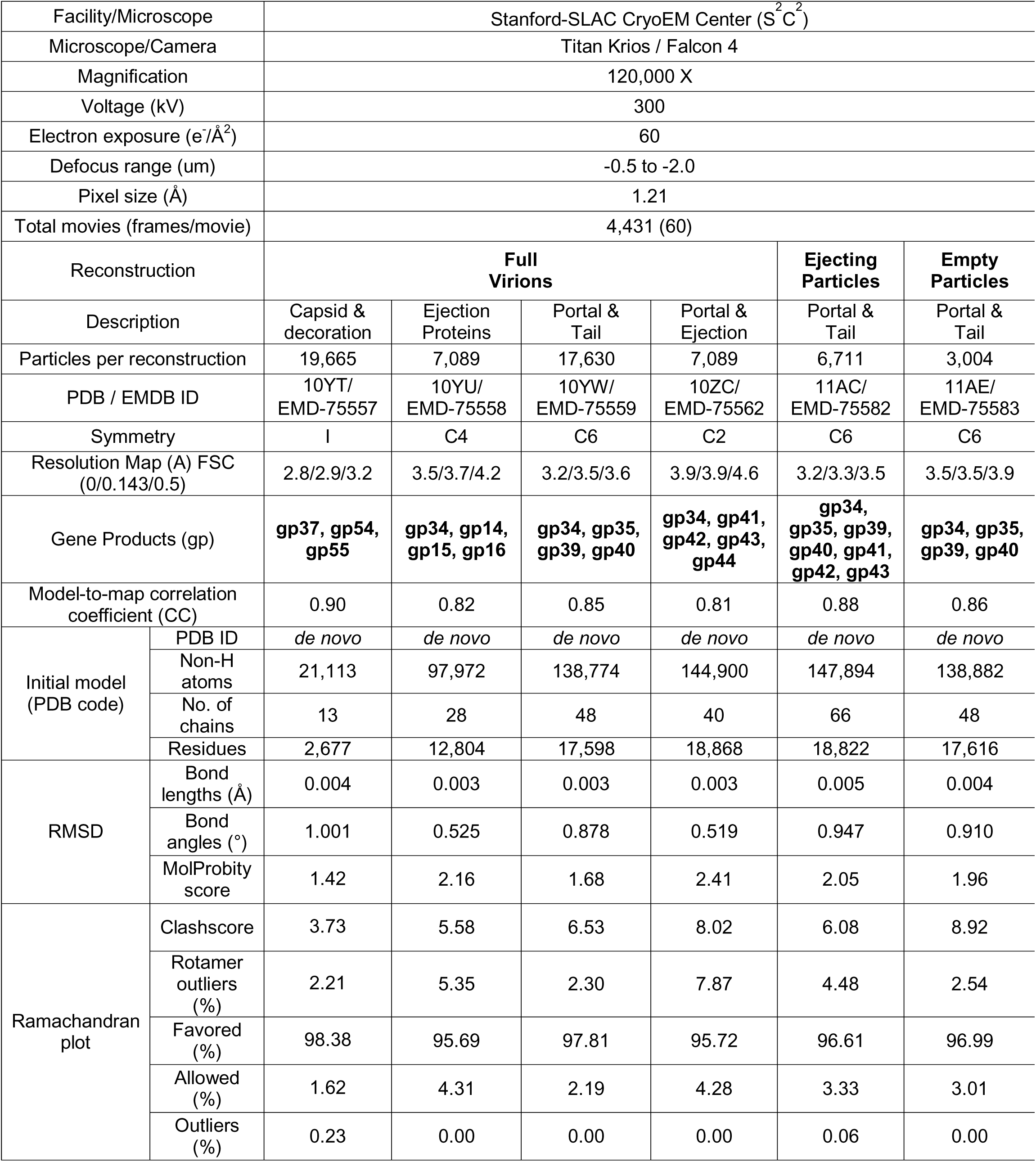
Cryo-EM models refinement statistics.

**Table 2.** Identification of Ar-KM Major Structural Proteins by LC-MS/MS.

| Protein | Mascot Score<sup>a</sup> | MW (Da) | Annotation |
| --- | --- | --- | --- |
| gp37 | 57872 | 37695 | Capsid protein |
| gp43 | 11175 | 143951 | Ejection protein (IMC) |
| gp55 | 10493 | 6904 | Decoration protein |
| gp42 | 8927 | 98191 | Ejection protein (PT) |
| gp35 | 8071 | 56088 | Portal protein |
| gp54 | 5693 | 10514 | Capsid spike |
| gp40 | 5580 | 91871 | Nozzle |
| gp34 | 5460 | 10465 | gp7.3-like |
| gp44 | 3841 | 28293 | Tail Fiber |
| gp45 | 2791 | 16325 | (putative) Tail Fiber 2 |
| gp41 | 2745 | 18855 | Ejection protein (OMC) |
| gp46 | 2305 | 33278 | (putative) Tail Fiber 3 |
| gp39 | 1919 | 21152 | Head to Tail-adapter |
| gp47 | 998 | 22582 | (putative) Tail Fiber 4 |
| gp36 | 194 | 33095 | (putative) Scaffolding protein |

<sup>a</sup> Statistical measurement of how well the observed MS data matches the identified protein sequence; scores greater than 50 represent correct identifications and can reflect relative protein abundance within a sample.

Overall, Ar-KM’s structural atlas presented in this paper comprises fourteen structural proteins (Fig. 2), <u>four</u> forming the capsid: gp37 (capsid protein), gp54 (spike protein), gp55 (decoration protein), and gp35 (portal protein); <u>four</u> inside the capsid, gp41, gp42, gp43 (ejection proteins), and gp34 (gp7.3-like factor); and <u>six</u> in the short tail apparatus: gp39 (head-to-tail adapter), gp40 (nozzle), gp44 (main tail fiber), and gp45, gp46, gp47 (putative tail fibers); in addition to <u>three</u> proteins, gp41, gp42 and gp34, in a post-ejection conformation. For all proteins identified in this study, we also performed a comparative bioinformatic analysis of sequence conservation using a curated dataset of 60 Ar-KM–like phages, spanning genome identities from 78.8% to 92.2% (Table S1). This analysis revealed striking patterns of conservation and divergence, providing insights into the biology and evolution of phiKMV-like phages.

### Architecture of *Pseudomonas* phage Ar-KM T = 7 capsid

Ar-KM has an icosahedral T = 7 capsid with an outer diameter of ∼650 Å (Fig. 3A). We reconstructed the Ar-KM icosahedral asymmetric unit at 2.8 Å resolution, which revealed the structure of the major capsid protein (MCP) gp37, spike protein gp54, and decoration protein gp55 (Fig. 3B), both real-space refined to a map-to-model CC of 0.89 (Table 1). Gp37 adopts the canonical HK97 fold [34], with clearly resolved domains comprising the E-loop (res. 52–89), P-domain (res. 90–140; 280–316), N-arm (res. 1– 33), and A-domain (res. 141–221, 230–279, 317–334) (Fig. 2). As is typical for T = 7 lattices, gp37 adopts two quasi-equivalent conformations in hexons and pentons to accommodate local curvature [35] (Fig. 3B). The closest match for the Ar-KM capsid protein is the *Shigella* phage Buco [36] (RMSD ∼ 2.7 Å), whereas phage T7 is only remotely related (RMSD ∼ 15.6 Å) [17]. Additionally, decoration (gp55) and capsid spike (gp54) proteins are present on the Ar-KM capsid surface (Fig. 2), also in analogy to phage Buco’s dense surface decoration gp47 and gp48 [36]. Two gp55 dimers per ASU are resolved at interfaces bridging neighboring gp37 subunits (across hexon–hexon and hexon–penton boundaries) (Fig. 3B). These dimers, which assemble into a four-helix bundle (Fig. 3C), act as structural staples, clamping adjacent capsomers and plausibly contributing to capsid stabilization and stress dispersion during assembly and maturation (Fig. 3B) [37]. Evolutionarily, most *Phikmvviruses* have a conserved gp55 (>85% identical) with just four phages (e.g., JB10, vB_PaeP_PE3, LUZ19, and vB_PaPhi_Mx1) where the identity drops to around 50% (Table S1), likely still consistent with the formation of a four-helix bundle.

**Figure 3.**
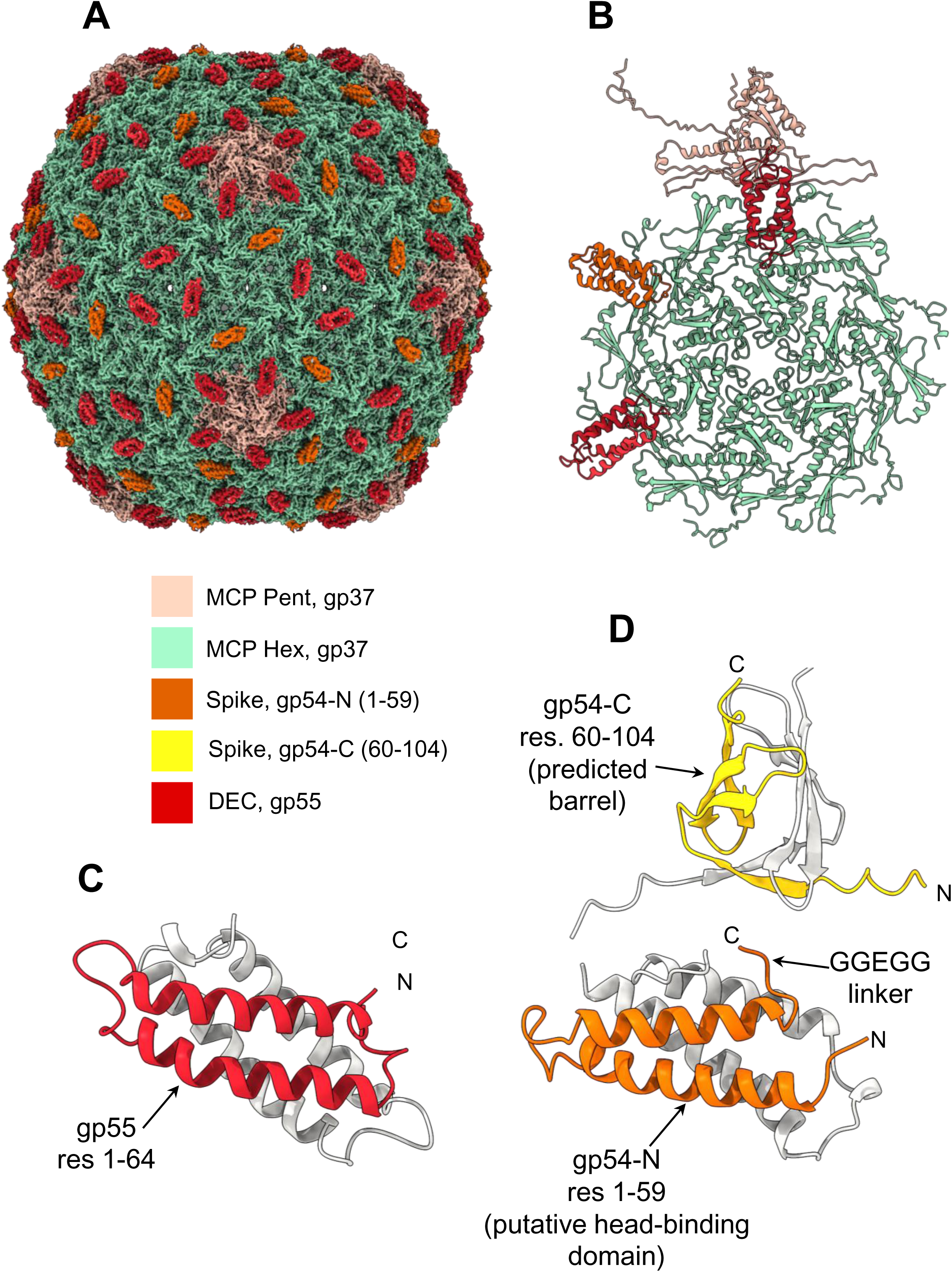
Reconstruction of Ar-KM capsid. (A) Icosahedral reconstruction of the Ar-KM T = 7 capsid determined at 2.8 Å resolution. (B) Ribbon diagram of the Ar-KM icosahedral asymmetric unit formed by seven copies of the MCP gp37, decorated with four copies of the decoration protein gp55 (red) assembled as homodimers and two spike protein gp54 (orange). (C) Ribbon representation of the putative decoration protein gp55 dimer, with one protomer shown in red and the other in gray. (D) Ribbon representation of the putative spike protein gp54 dimer, with one subunit shown in orange and the other in gray. The flexible spike domain predicted by AlphaFold 3 is highlighted in yellow (one protomer) and gray (second protomer).

Although gp54 was readily detected by LC–MS/MS in purified particles, it is partially smeared in both icosahedral and C5-maps, precluding the generation of a complete atomic model. Interestingly, gp54 comprises two domains: residues 1–59 form a helical core linked by a flexible GGEGG motif to a C-terminal domain (res. 60-104) that projects outward toward the surrounding milieu. This region, smeared in the reconstruction, is predicted by AlphaFold 3 to fold into a small barrel (Fig. 3D). The N-terminal cores of gp54 and gp55 are highly similar, with near-identical structures (RMSD ∼0.426 Å over the first 55 residues) (Fig. 3C, D). Evolutionarily, gp54 is broadly conserved among phiKMV-like phages, with 52 of 60 homologs sharing >95.2% sequence identity (Table S1). However, a handful of phages (e.g., vB_PaeP_PE3, Ab05, PJNP013, DL62, BARC01, and vB_PaePA10145phi1_HR2) encode variants with markedly divergent C-termini. Within the modeled N-terminal fragment of gp54 (residues 1–59), all residues involved in dimerization and capsid interaction are strictly conserved. Notably, sequence divergence among these homologs begins at residue 59, supporting the idea that residues 1–59 define a putative head-binding domain that anchors gp54 while displaying the variable C-terminus outward (Fig. 3D).

### Ar-KM short tail apparatus

We computed a localized reconstruction of the Ar-KM tail apparatus from FV that was then symmetrized using C6 symmetry, achieving a maximum resolution of 3.5 Å (Table 1, Fig. 4A). This density (Fig. S4) enabled identification and *de novo* modeling of the portal protein gp35, the head-to-tail adapter (HT-adapter) gp39, the nozzle (gp40), and the trimeric tail fiber (gp44) (Fig. 2) that together were refined to a CC of 0.85 at 3.5 Å resolution (Fig. 4B, Table 1). Evolutionarily, these proteins are nearly invariant across the 60 *Phikmvviruses* (>99% identical) analyzed in this study (Table S1). Overall, the mature virion tail extends approximately 50 Å into the capsid interior and 100 Å outward, beyond the capsid surface. The Ar-KM portal protein was modeled using a C6 map, which unexpectedly showed higher-quality density than the C12 map, underscoring the lack of perfect C12 symmetry across all subunits (see below). Ar-KM adopts the classical portal fold, comprising barrel, wing, stem, and crown domains (Fig. 2) [37]. It is most similar to the portal from the *Shigella* bacteriophage HRP29 (PDB: 8ES4), and shares approximately 29% sequence identity with the T7 portal protein. The lower portal crown exhibits near-perfect C12 symmetry, while the barrel α-helices deviate from ideal C6 symmetry, as described in the following section. Attached to the bottom of the portal is the HT-adapter gp39 (Fig. 4B), which is structurally similar to Cyanophage P-SCSP1u [38]. It features a helical core with a C-terminal α−helix that inserts into the portal protein interface, similar to other podophages [25, 26]. However, the C-terminal core also contains an insertion domain spanning residues 44-110 (Fig. 2) that makes contact with the N-terminus of the tail fiber gp44. Below the HT-adapter, we observed six copies of the nozzle protein gp40 (Fig. 4B), which in Ar-KM consists of a double-interrupted β-propeller (Fig. 2) comprising three domains: a central 6-bladed β-propeller, an N-terminal insertion (NTI) domain emanating from the second blade (res. 131-349) that points outward and forms the nozzle tip, and a C-terminal insertion (CTI) domain (res. 619-824) that assembles onto the HT-adapter dodecamer.

**Figure 4.**
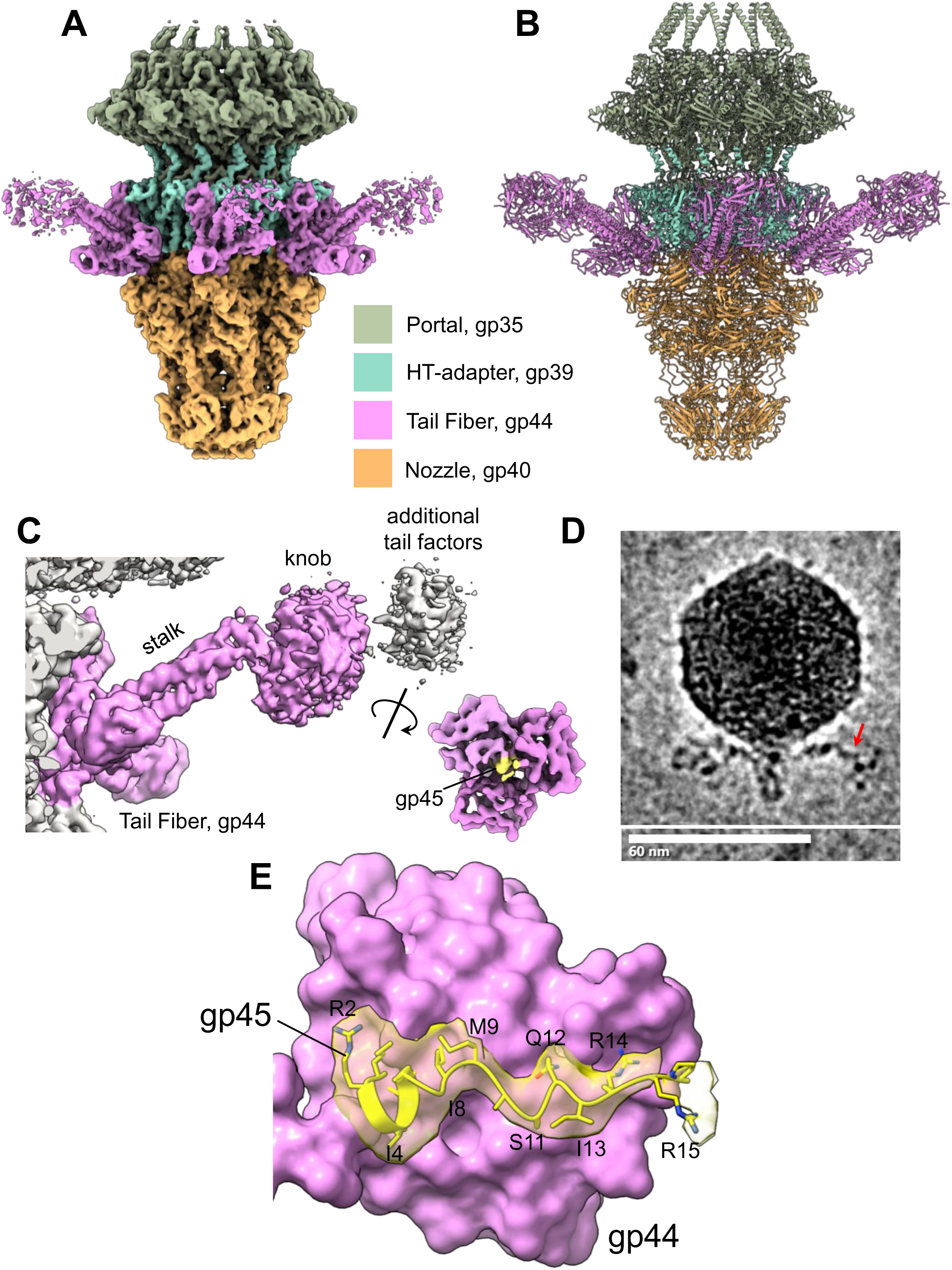
Reconstruction of Ar-KM tail and tail fiber. (A) The 3.5 Å reconstruction of Ar-KM tail apparatus from FV is color-coded by subunits as shown in Fig 2. (B) Ribbon diagrams of all the atomic models built *de novo* within the cryo-EM density in panel (A). (C) C6-averaged focused reconstruction of the tail fiber gp44 (magenta) from FV reveals additional spurious density juxtaposed to the gp44 knob. A rotated view of the knob reveals density (in yellow) at the center of the knob trimeric interface. (D) Representative cryo-micrograph of Ar-KM mature virion showing density emanating from the tail fiber (red arrow). (E) 3.9 Å resolution, C1-averaged density (transparent yellow) overlaid on the N-terminus of gp45 (residues 2-15). In the background, shown as a pink solvent surface, is one of the three subunits of the gp44 knob.

### Ar-KM tail fibers anchor a flexible molecular complex

The main tail fiber anchor attached to the Ar-KM tail is encoded by the gene *gp44*, which produces a ∼130 Å trimeric fiber that extends outward (Fig. 4A, B). This gene is conserved, with an average sequence identity of 97.2% across the 60 *Phikmvviruses* listed in Table S1. Six gp44 fibers decorate the Ar-KM tail (Fig. 4A, B) and attach at the interface between the HT-adapter and the nozzle, primarily stabilized by the HT-adapter insertion domain (Fig. 2). However, the fibers are shorter than the phage nozzle and do not extend beyond the capsid, making them unlikely to serve as effective attachment points for the pilus. We modeled the entire sequence of the tail fiber (Figs. 2 and 4C), which is rather short and stocky, consisting of an N-terminal six-stranded β-barrel (res. 1-84) shaped like a petal, an α-helical coiled-coil (res. 85-142), and a C-terminal knob (res. 143-251), somewhat similar to that of phage Sf6 [26, 39]. The knob is flexibly connected to the coiled coil, which we speculate can pivot relative to the coiled-coil stalk, as also indicated by the partially smeared density (Fig. 4A).

In the reconstruction, gp44 is also oriented upward, which seems incompatible with a direct role in pilus binding. This prompted us to carefully review the density near the knob and search for additional, loosely associated densities. Inspection of raw FV micrographs revealed density extending from the tail fiber knob (Fig. 4D), which was poorly resolved in the C6 map (Fig. 4C). We calculated a focused reconstruction of the tail fiber using a tight mask, followed by threefold averaging to enhance the signal-to-noise ratio. This map revealed a distinct density within the trimeric tail fiber knob that corresponds to the N-terminus of gp45 (Fig. 4E), an ORF adjacent to the tail fiber gene (Fig. 2) detected in our preparation by MS analysis (Table 2). The N-terminus of gp45 forms a short and hydrophobic α-helix (2-RG**II**AG-7) that inserts into a deep crevice within the trimeric knob, creating an unusual 3:1 symmetry mismatch. Gp45 residues C-terminal of the short helix (8-**IM**ASQ**I**RR-15) adopt an extended conformation with visible side chains contacting the lumen of the trimeric knob. No discernible density is available for the other putative subunits of the tail fiber (gp45 aa16-237, gp46, and gp47). It is possible that the three ORFs located downstream of the tail fiber gene, gp45, gp46, and gp47, contribute to the filamentous density observed in the micrographs. Evolutionarily, these gene products are significantly more divergent than other structural proteins in Ar-KM (Table S1). Gp45 exhibits an average sequence identity of 66.80%, but this average hides wide variability, with sequence identity ranging from 26.0% to 95.4% across different phiKMV-like phages. Similarly, gp46 (average sequence identity 73.14%, range 22.3–99.7%) and gp47 (average sequence identity 81.68%, range 47.5–99.5%) show markedly lower conservation within the 60 phiKMV-like phages analyzed in this study. This elevated sequence variability is consistent with a potential role in tail fiber-associated functions involved in host recognition.

### Organization of the ejection proteins before genome ejection

The localized reconstruction of FV’s unique vertex showed an ordered density above the portal protein, which we model as the three ejection proteins gp41, gp42, and gp43 (Fig. 2), similar to those of phage T7 [19]. The density was most clearly visible in a C4-averaged localized reconstruction, which revealed an assembly 160 × 210 Å wide that covers the portal protein and extends into the capsid (Fig. 5A, B). Interestingly, a cutaway view of the ejection assembly revealed an inner channel approximately 30 Å in diameter, co-axial with the portal channel. Furthermore, we identified a new density (cyan in Fig. 5B), that we assigned to gp34 (Fig. 2). Overall, we built atomic models for four copies of the gp16-like factor gp43 (Fig. 5C), eight copies of the gp15-like factor gp42 (Fig. 5D), eight copies of the gp14-like factor gp41 (Fig. 5E), and eight copies of the gp7.3-like factor gp34 (Fig. 5F). Overall, the ejection proteins and gp34 were refined to a CC = 0.81 against a 3.6 Å density (Table 1) and present a 4:8:8 stoichiometry (gp43:gp42:gp41) with an additional eight copies of gp34 bound primarily to gp41.

**Figure 5.**
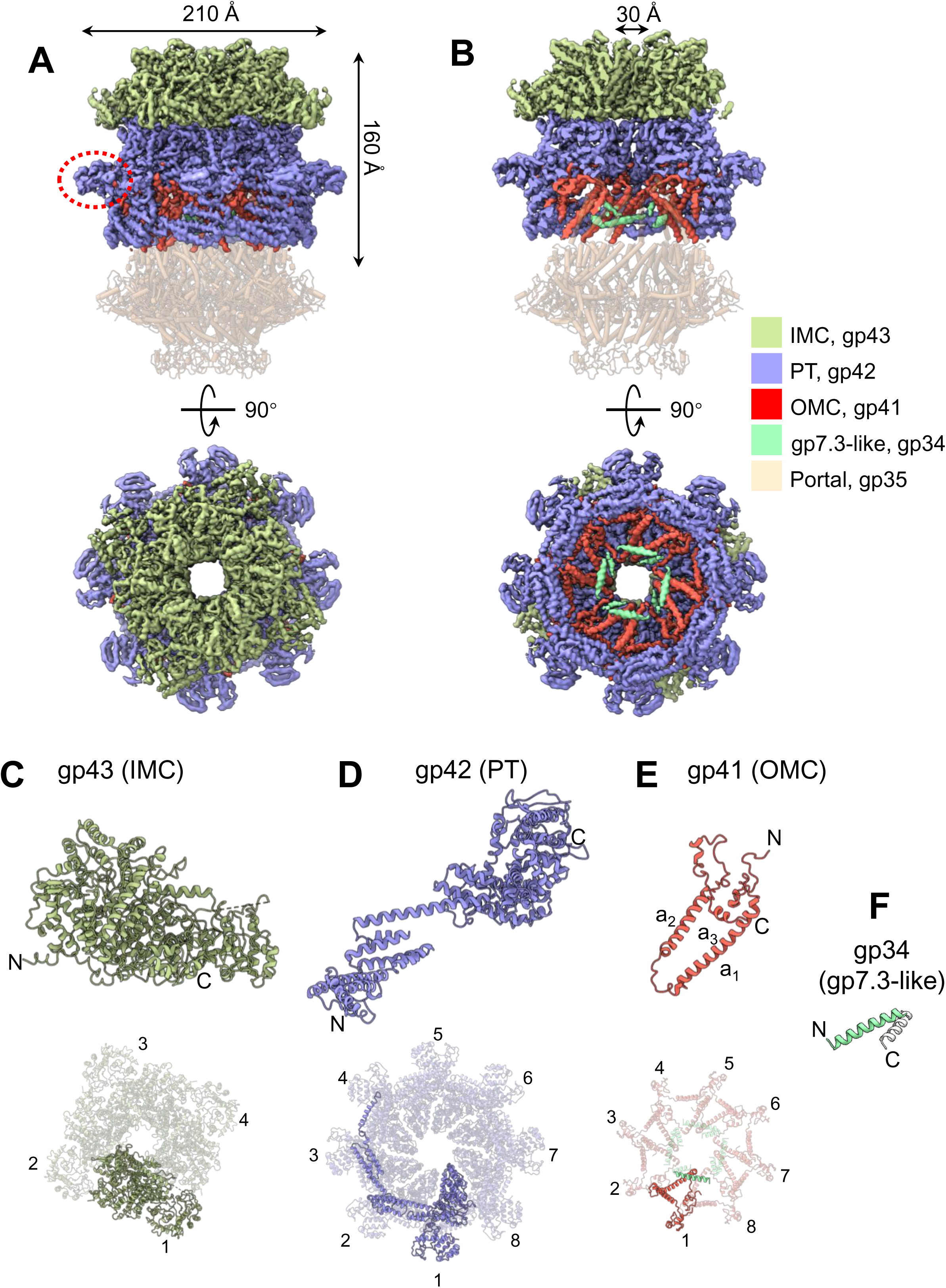
Ar-KM ejection proteins form coaxial rings stacked on the portal protein. (A) Side view of the experimental density for Ar-KM ejection proteins, extracted from the C4-symmetry reconstruction of FV at 3.6 Å resolution. The density is overlaid on the portal protein, shown as a semi-transparent ribbon diagram with cylindrical helices. The region of gp42 circled in red indicates the T4 lysozyme-like domain. (B) Cutaway view of the density in (A), revealing a ∼30 Å lumen within the ejection protein stack. (C-E) Ribbon diagrams of Ar-KM ejection proteins gp43 (green), gp42 (blue), and gp43 (red). Each ejection protein is shown as a protomer (top) and in the oligomeric state observed in FV (bottom). (F) The ribbon diagram of gp34, visible as two α-helices shown in green and gray, is associated with gp41 at the interface with the portal protein.

Bioinformatic analysis suggests that Ar-KM gp41, gp42, and gp43 are all conserved, with sequence identity exceeding 91.2%, 93.3%, and 90.7%, respectively, within phiKMV-like phages (Table S1) and closely resemble the ejection proteins of phage T7 [17]. Foldseek identified a lysozyme domain in gp42 residues 740-881 (red circle in Fig 5A and Fig. S5A, B), which superimposes with RMSD 3.8 Å to the T4 lysozyme (gp5) [40, 41]. Perhaps more surprising is the position and stoichiometry of gp34 (Fig. 5F). This gene product is more than 96.9% identical across *Phikmvviruses* (Table S1), underscoring its vital function. AlphaFold 3 predicts gp34 as a continuous α-helical polypeptide, but only two partial segments could be modeled in the C4 reconstruction at our current resolution (∼3.6 Å) (Fig. 5E). Where resolved, two gp34 helices adopt an inter-helix angle of ∼75°, positioned inside the gp42:gp41 cage and close to the center (Fig. 5E, F). Focused 3D classification along the gp34 N and C-terminal extensions reveals multiple alternative densities; however, particle scarcity and pronounced heterogeneity preclude a coherent atomic path for these distal segments. Interestingly, the assembly of gp34 and gp41 onto the portal protein barrel disrupts the C12 symmetry of the barrel helices. The C6 (and C12) reconstruction reveals that while the portal crown is strictly C12, the barrel helices cluster into four trimers, each making contact with two copies of gp41 and two gp34 (Fig. S6A). Thus, the symmetry 12:8 mismatch between the portal and gp41 is reconciled by a local conformational change in the portal barrel that adopts a quasi-4-fold symmetric structure. Each barrel helix deviates from its expected position in a C12 ring around 2.5 Å (Fig. S6B).

### EJP tail reveals the basis for genome gating

The second population of phage particles reconstructed in this study, EJP (Fig. 1C), has a fully open tail, with a continuous lumen that varies in diameter between 30 and 50 Å, at all points large enough to accommodate double-stranded DNA (dsDNA) (Fig. 6A). Inspection of the 3.3 Å C6 tail map revealed density lining the nozzle channel that we interpreted as the ejection proteins gp41 and gp42, as well as gp34 (Fig. 6B). Specifically, we modelled residues 12–153 of gp41 (of 180 residues), residues 5–35 of gp42 (of 898 residues), and a short segment of gp34 comprising residues 34–81 (of 98 residues). The three proteins form a heterotrimeric complex (Fig. 6B) that is repeated six times along the nozzle lumen (Fig. 6B). The atomic models of gp41:gp42:gp34 were refined to a CC = 0.88 at 3.3 Å resolution (Table 1).

**Figure 6.**
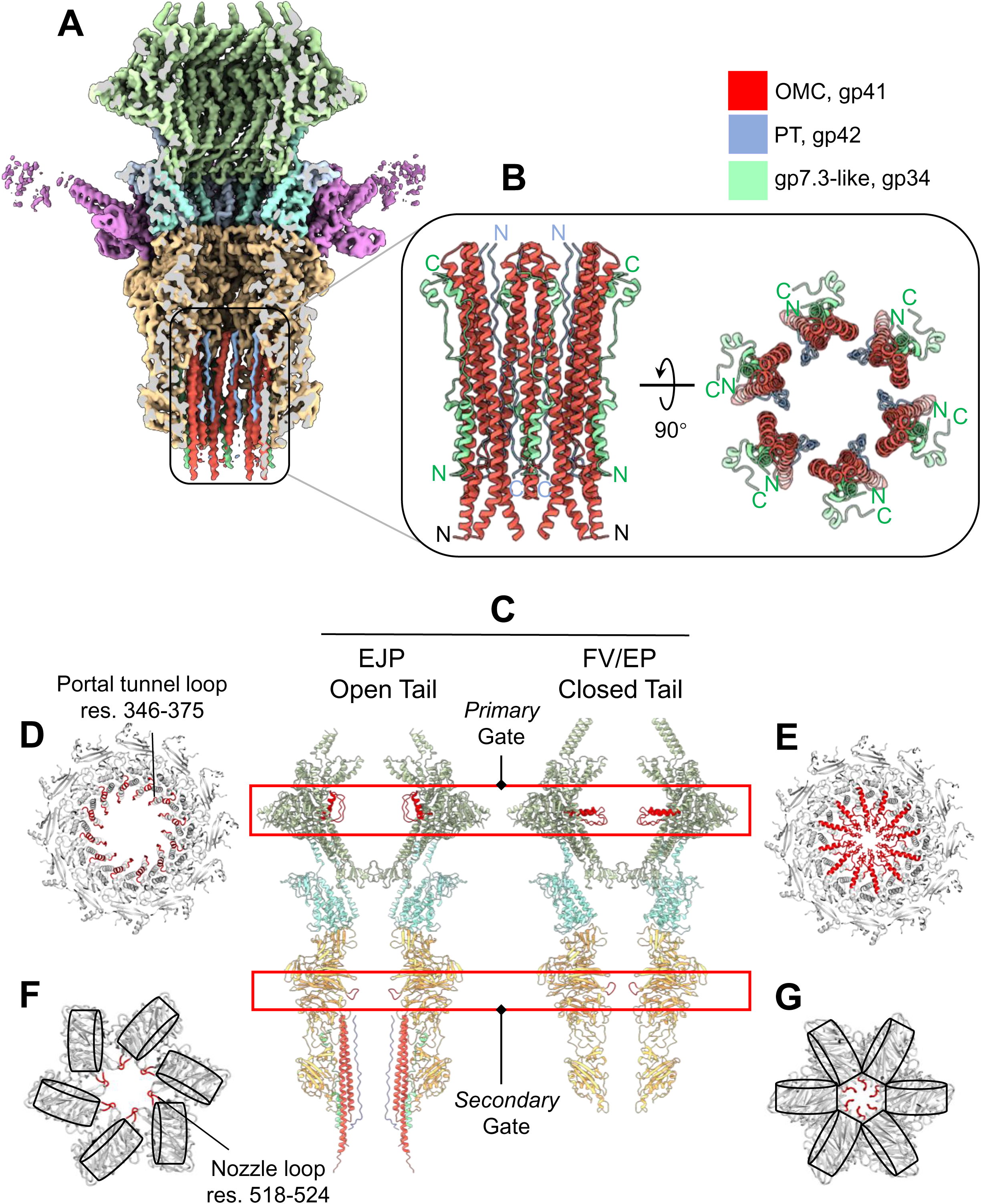
Ar-KM ejecting particles reveal the mechanisms of genome gating. (A) A cutaway view of the C6 cryo-EM reconstruction of the Ar-KM tail apparatus from EJP. The 3.3 Å experimental map is color-coded by subunit, as in Figure 4A. (B) Ribbon diagram of the heterotrimeric complex comprising gp41 (red), gp42 (light blue), and gp34 (cyan), present in six copies lining the nozzle lumen. (C) Cutaway views of the EJP open tail (left) and the FV closed tail (right). Red insets highlight the primary gate in the portal protein and the secondary gate in the nozzle. (D) Top view of the EJP portal protein, sliced to reveal the primary gate. Residues 346-375 are colored red to emphasize the open channel. (E) Top view of the FV portal protein, sliced to reveal the primary gate. Residues 346-375 are colored red to emphasize the closed channel. (F) Top view of the EJP nozzle, sliced to reveal the secondary gate. Residues 518-524 are colored red to emphasize the open nozzle lumen. (G) Top view of the FV nozzle, sliced to reveal the secondary gate. Residues 518-524 are colored red to highlight the closed-nozzle lumen. In panels (F) and (G), schematic cylinders are overlaid on the β-propeller domains of gp40 to illustrate the relative organization of nozzle subunits.

EJP tail lumen is fully open, generating a continuous aqueous channel from the capsid to the outside, with an internal diameter of 30-50 Å (Fig. 6C). In contrast, FV and EP have two major gating points: the portal primary gate and the nozzle secondary gate (Fig. 6C), similar to phage T7 [42] or the cyanophage P-SCSP1u [38]. Comparing EJP and FV enabled us to detail the organization of the two gates. The primary gate is at the portal protein, gp35, which adopts two dramatically distinct conformations in EJP and FV (RMSD 3.8 Å). The primary gate is generated by the retraction of the portal helix-loop comprising residues 346-375, which are collapsed laterally in EJP (Fig. 6D), but are swung into the lumen of FV and EP, restricting the lumen diameter to ∼6 Å (Fig. 6E). The secondary gate is within the nozzle, gp40, which adopts two distinct conformations in the open tail of EJP versus closed FV and EP (Fig. 6F, G). In the open tail, the nozzle subunits adopt a loose organization with the six β-propellers slightly skewed relative to each other (Fig. 6F), featuring a lumen approximately 30 Å wide, with a loss of subunit interaction at the level of the NTI-domains (Fig. 2), as described in the next section. In contrast, the closed FV tail (Fig. 6G) has β-propellers in a tense quaternary organization, touching each other laterally and projecting an internal loop (res. 518-524) into the lumen, which restricts the channel to sub-10 Å. A global superimposition of the nozzle subunits, arranged as hexamers, results in an RMSD of 3.3 Å. The main variations are in the NTI-domain insertion, while the CTI-domain and β-propeller show minimal changes in the open and closed tails. In contrast, the NTI-domain undergoes dramatic conformational change, showing a maximum displacement of ∼10 Å between the two states (Fig. 7A).

**Figure 7.**
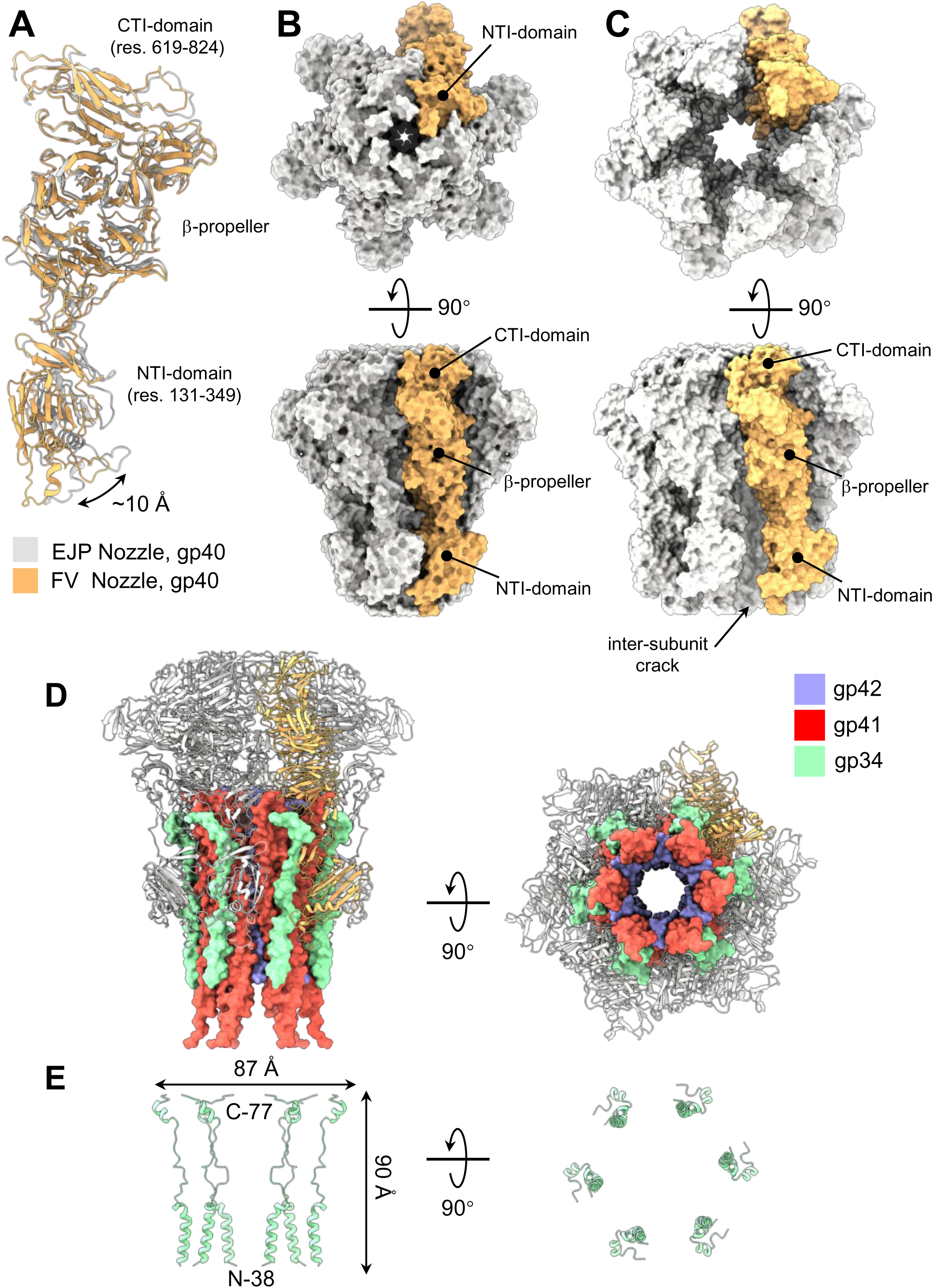
Gp34 stabilizes the open nozzle. (A) Overlay of ribbon representations of the FV and EJP nozzles, showing an approximately 10 Å displacement of the NTI around the β-propeller domain. (B) Surface representation of the closed FV nozzle in bottom (top) and side (bottom) views, with one protomer colored orange and the remaining five in gray. (C) Surface representation of the open EJP nozzle in bottom (top) and side (bottom) views. One protomer is colored orange, and the remaining five are shown in gray. (D) Ribbon representation of the open EJP nozzle in bottom (right) and side (left) views. One protomer is colored orange, and the remaining five are shown in gray. All six copies of gp41 (red), gp42 (blue), and gp34 (cyan) lining the lumen are displayed as solvent surfaces. (E) Ribbon diagram of the six gp34 subunits seen inside the EJP nozzle in side (left) and top (right) view.

To understand how the nozzle opens in EJP, we analyzed the quaternary structure of gp40 in open and closed states. In the closed tail (Fig. 7B), six gp40 subunits stack laterally, sharing an extensive binding interface that spans the three domains, from CTI, at the boundary with HT-adapter, to the distal nozzle tip formed by NTI-domains. No solvent channel is observed between gp40 subunits, due to the tight subunit interface that is stabilized by 20 hydrogen bonds, three salt bridges, and 268 non-bonded contacts. However, in EJP (Fig. 7C), the hexameric assembly of gp41:gp34 expands the lower part of the nozzle cavity, breaking half of the gp40 lateral interface that shows a visible crack from the outside. This opening results from two factors. On one side, gp34 adheres to the gp40 NTI-domain like a wedge (Fig. 7D), forming a cage within the nozzle (Fig. 7E) that prevents lateral association of gp40-NTIs, as seen in FV, and provides a contact surface for gp41. On the other hand, the ejection protein gp41 inserts rigidly into the disrupted NTI:NTI interface, folding like a helical hairpin (Fig. 7D). In contrast to gp41 and gp34, the short gp42 fragment visible in the EJP reconstruction lines the open nozzle channel and makes no contact with the nozzle subunits (Fig. 7D). Together, gp34 and gp41 stabilize the open-gate structure of EJP, keeping the lumen unobstructed.

### Bioinformatic identification of gp34 (ejection wedge) orthologs

A cursory examination of the genomes of one member from each genus within the order *Autographivirales* (205 genomes, Table S2), covering phages infecting organisms as different as *E. coli*, *Pseudomonas*, *Rhyzobium,* or *Synechococcus*, shows that the portal genes (which are sufficiently conserved to be identified via domain conservation) are always preceded by at least one small ORF. There is insufficient sequence conservation among most of these ORFs to assign function by homology, but most of these ORFs share four features with Ar-KM gp34 (Fig. 8A) and T7 gp7.3 (Fig. 8B), leading us to suggest they could be functional orthologs. <u>First</u>, they are short, typically between 100-140 amino acids (although they can be as short as 45aa and as large as 450aa); second, they are predicted to be highly helical; <u>third</u>, both gp7.3 and gp34 possess N-and C-terminal segments that are positively charged (R, K); <u>fourth</u>, the central section is low complexity, richer in Proline, Alanine and Glutamine on the N-terminal side and negatively charged amino acids on the C-terminal side (Fig. 8A). Notably, our structure indicated that the N-terminal helix visible in EJP encompasses residues 38-54, point downward (Fig. 7E), toward the OM, while the second helix (res. 70-77) points inside the lumen. Although we do not see residues from the C-terminal basic cluster, it is possible that these residues would interact with the DNA inside the capsid. These observations suggest that reliance on wedge proteins to open the nozzle during DNA ejection may be a conserved feature of Autographivirales and warrant further studies of less-studied phages to confirm this hypothesis.

**Figure 8:**
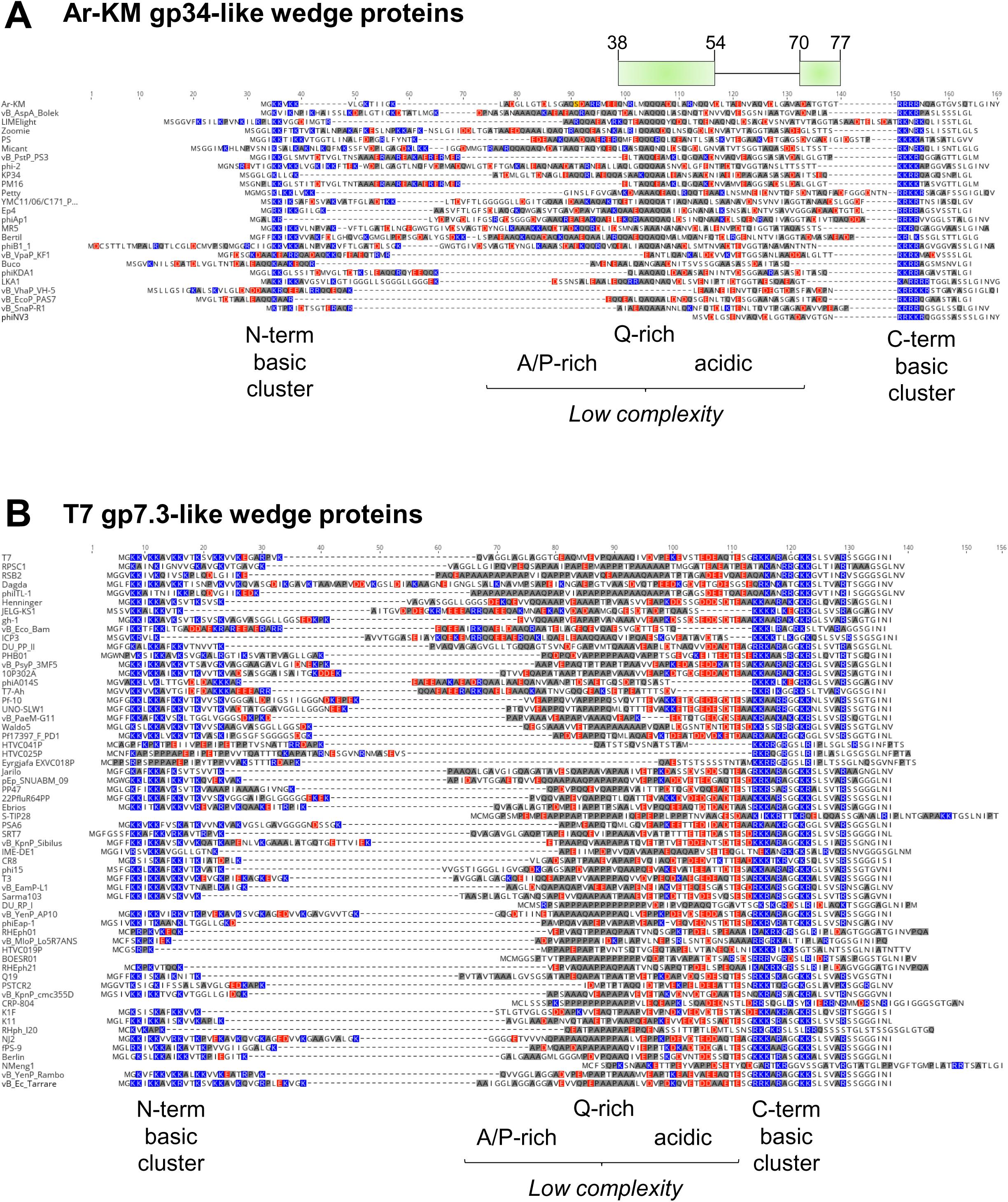
Ejection wedge orthologs show a signature motif. ORFs located upstream of the portal gene across diverse Autographivirales genomes were aligned. (A) Ar-KM–like gp34 homologs were extracted, and alignments were manually curated to highlight characteristic sequence features: enrichment of basic residues (blue) at both termini, a central low-complexity region enriched in alanine, glutamine, and proline (A/Q/P; dark grey), and a strongly acidic stretch separating the AQP-rich core from the C-terminal basic cluster. (B) Proteins homologous to gp7.3 were identified in phiKMV-like phages. These sequences display a similar sequence pattern, characterized by two basic clusters flanking a central low-complexity region.

### Post-ejection conformation of gp41 inside the nozzle of EJP

Comparing gp41 from FV and EJP offers a comprehensive view of the conformational changes that happen during genome ejection. Before ejection, gp41’s two main α-helices (Fig. 9A) are connected by a five-residue non-helical turn that inverts the helix trajectory; together, eight gp41 copies form a flat oligomeric assembly approximately 155 Å wide, barely 70 Å in height, decorated by gp34 on the interior (Fig. 7D). In contrast, in EJP (Fig. 9B), gp41 forms a continuous a-helical hairpin, whereby both helices a1 (res 42-82) and α2 (res. 94-120) are twice as long as before ejection, due to the helical folding of nearby amino acids. Six gp42 hairpins line the inner nozzle walls, maintaining alternating contacts with gp34 and gp40 NTI (Fig. 7D). Additionally, gp41 oligomerization stoichiometry changes from eight copies to six, which expands the inner lumen diameter below the secondary gate from about 22 Å pre-ejection to a 30 Å after ejection (Fig. 9A, B). Inside the gp41 tube, gp42 N-terminal residues 5-35 line the walls (Fig. 7C). gp42 points its entire C-terminus to the outside, and gp41 projects both its N-and C-termini to the outside of the nozzle. The two proteins, however, form an extensive binding interface within the nozzle, involving 12 hydrogen bonds, indicating that they remain associated during ejection.

**Figure 9.**
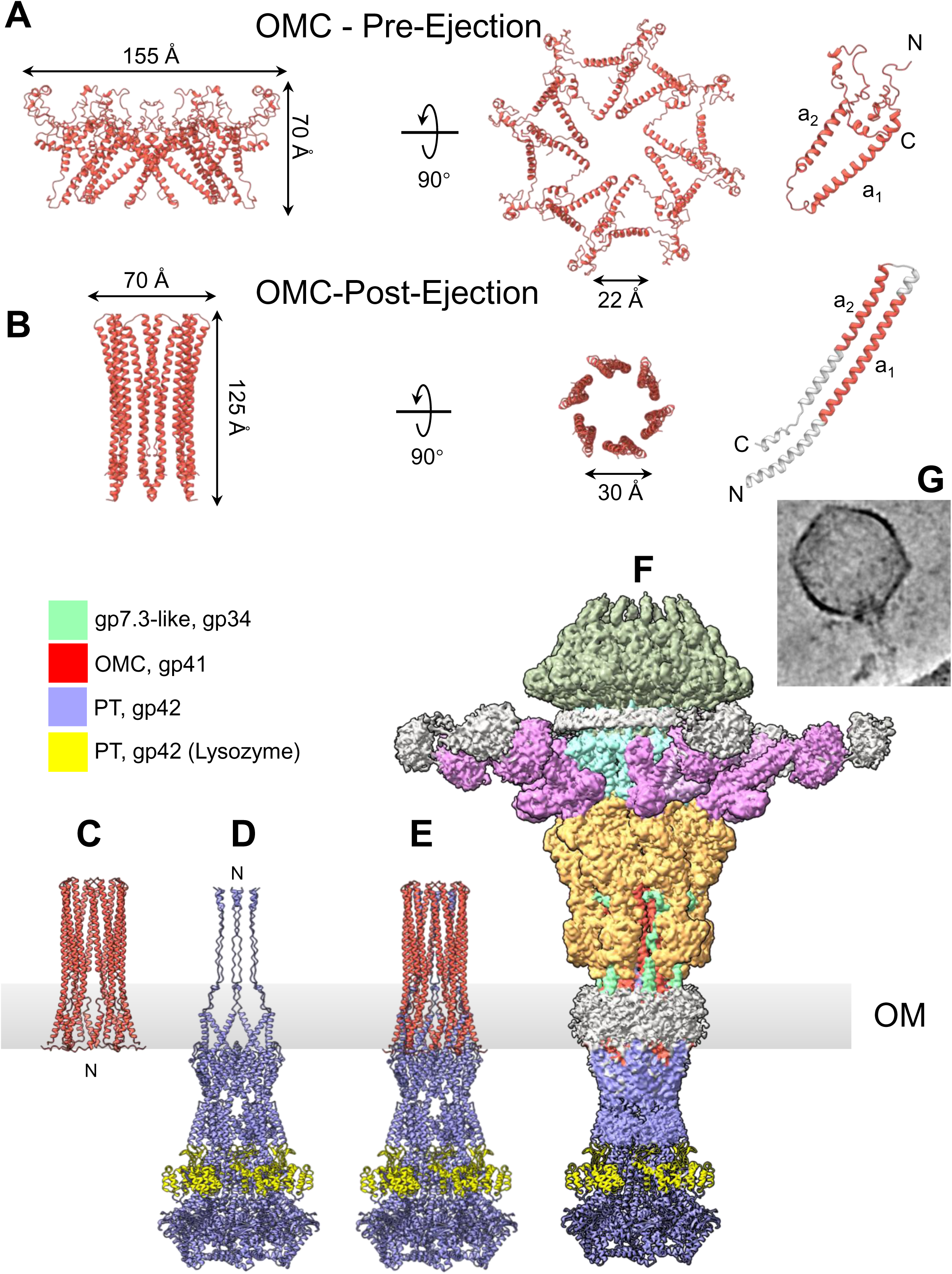
Post-ejection conformations of Ar-KM ejection proteins gp41 and gp42. (A) Pre-ejection conformation of the ejection protein gp41 from FV. The left and center panels show two rotated views of the oligomer, while the gp41 protomer is shown on the right. (B) Post-ejection conformation of the ejection protein gp41 from EJP. The left and center panels show two rotated views of the oligomer, while the gp41 protomer is shown on the right. Regions of gp41 that adopt a helical fold in the post-ejection conformation are shown in gray. (C–E) AlphaFold 3 models of hexameric gp41 (red, in (C)), hexameric gp42 (blue, in (D)), and the gp41:gp42 complex (red and blue, in (E)). (F) Low-resolution cryo-EM reconstruction of a subset of 1,725 EJP in which the nozzle is extended by a density that resembles that of the AlpaFold 3 model of gp41:gp42, which is overlaid on the low-resolution density. (G) Representative cryo-micrograph of Ar-KM from the subset of 1,725 particles with the extended nozzle density. In panels (C–F), the light gray box indicates the putative OM.

The partial structures of gp41 and gp42 observed in EJP prompted us to model their assembly after ejection using AlphaFold 3 [43]. Using the number of subunits observed in EJP, equal to six subunits, we predicted a hexameric assembly of gp41 (Fig. 9C), which is similar, but more elongated than the experimentally observed structure (Fig. 9B). The N-terminal residues 1-25 and C-terminal 151-181 of gp41 contain a predicted membrane spanning helix and are not visible in EJP reconstruction. Similarly, using a stoichiometry of six, we predicted a structure of gp42 assembled like a PT (Fig. 9D). Notably, AlphaFold 3 mostly predicted the gp42 N-termini as random coiled, and, as seen in EJP, the gp41:gp42 assembly has the gp42 N-termini lining the lumen formed by gp41 helices (Fig. 9E). This model was supported by a low-resolution reconstruction that featured an extended nozzle (Fig. 9F). This reconstruction was achieved by applying a cylindrical mask along the nozzle, followed by an additional round of 3D classification, which isolated a subset of 1,725 particles. After heterogeneous refinement, this subset displayed a clearly defined axial conduit through the nozzle. Since no membrane mimetic was introduced and the micrographs (Fig. 9G) show no particle–particle contact (e.g., virions are not adjacent), phage-to-phage triggering is unlikely. Therefore, we hypothesize that air–water interfaces or grid–hole boundaries can serve as *in situ* triggers for ejection in vitrified samples and may do so more consistently than inter-phage contacts under our conditions.

### Peptidoglycan hydrolase activity associated with ejection proteins

In the predicted quaternary structure of gp42 PT, the lysozyme domains face outward, as shown in Fig. 9D (yellow). In T7, the PT protein (gp15) lacks a lysozyme domain, which instead is part of the IMC protein gp16. The lysozyme domain is responsible for the partial degradation of the cell wall, allowing the PT and IMC to reach the IM [44]. The visual similarity between the predicted transmembrane ejectosome of Ar-KM and that of T7 is remarkable (Fig. S5C). The two assemblies are similar in length and position the transglycosylase domain at comparable heights, likely corresponding to the cell wall location. To assess the evolutionary distribution of this feature, we performed a bioinformatic analysis of the order *Autographivirales*, which includes phiKMV-like phages. Among 205 genomes analyzed, 127 encode identifiable lysozyme/transglycosylase domains within predicted ejectosome proteins (Table S2). These domains occur in three distinct configurations: (i) at the N-terminus of the IMC (first ∼160 residues), as in T7 (86/127; 67.7%), spanning diverse hosts including Enterobacteriaceae, Pseudomonadales, and marine phototroph-associated bacteria; (ii) at the C-terminus of the PT (∼last 300 residues), as observed in Ar-KM (31/127; 24.4%); and (iii) centrally within the PT, around residue ∼300, a configuration restricted to phages of the *Dunnvirinae* subfamily (10/127; 7.9%). Despite this variability, the combined length of the OMC+PT+IMC module remains constrained within a narrow range (2378 ± 162 amino acids), consistent with the requirement to span the relatively fixed width of the periplasmic space. Together, these findings support a model in which ejection proteins are evolutionarily plastic, undergoing domain fusion and reorganization, while preserving the overall topology and functional architecture of the ejectosome.

## DISCUSSION

In this study, we used an integrative structural and evolutionary approach to define the molecular architecture of phage Ar-KM, a therapeutic *P. aeruginosa* phage from Armata Pharmaceuticals’ collection. The serendipitous structural heterogeneity inherent in the Ar-KM preparation enabled us to visualize three distinct viral states, thereby allowing us to assemble a structural atlas of its constituent protein. We identified the ejection proteins gp41, gp42, and gp43 in their pre-ejection conformations and, for gp41 and gp42, we further resolved key features of their post-ejection states. We also characterized gp34, a highly conserved gene product within the phiKMV-like family, which remains intimately associated with gp41 both before and after ejection. Collectively, these findings illuminate three aspects of Ar-KM biology that are broadly generalizable to other podoviruses, including the model phage T7.

<u>First</u>, Ar-KM displays fully ordered ejection proteins arranged as a stacked assembly on the portal protein prior to ejection. A similar organization has been described for phage T7 [17, 18] and, more recently, for the *E. coli* phage E1004 [45]. Our work highlights the remarkable structural plasticity of the OMC protein gp41, which undergoes a dramatic conformational rearrangement upon genome release, forming a hexameric channel composed of a helical hairpin. Beyond providing a more complete and higher-resolution structural model of gp41 (the functional equivalent of gp14 in T7), we also delineate the interface between gp41 and the PT subunit gp42 (gp15 in T7). The manner in which these proteins assemble after ejection is functionally analogous to the insertion of the phage DEV PT N-termini into the OMC lumen [28], suggesting that both ejection-protein architecture and assembly mechanisms are conserved [20]. Although primary-sequence similarity among podophage ejection proteins is minimal, often below standard thresholds for bioinformatic detection, their higher-order assembly rules appear surprisingly well conserved. Consistent with this view, our structures reveal a reduction in subunit number from the pre-ejection octameric conformation of gp41 and gp42 to the hexameric state formed after ejection. Unlike earlier post-ejection structures derived from recombinantly purified proteins [21, 22, 46], which may reflect *in vitro* artifacts, the ejection protein structures captured in EJP in this study were identified *in situ*, reflecting an authentic stage of the phage life cycle and confirming the loss of two gp41 and gp42 subunits upon ejection. An analogous reduction in subunit stoichiometry has been observed in phage DEV, where the OMC protein gp73 and the PT protein gp72 transition from dodecameric assemblies in the mature virion to nonameric forms post-ejection [28]. A recent study suggests that a similar mechanism may operate in phage N4 [31]. As more ejection proteins are structurally resolved, a coherent picture is emerging in which the principles governing ejection-protein organization, membrane penetration, and post-ejection assembly are deeply conserved across diverse podoviruses. This observation further illustrates that phage evolution is more coherently captured by conserved structural principles than by rapidly evolving primary sequences [47].

<u>Second</u>, our study identified the Ar-KM genome-gating mechanism, which can be generalized to phiKMV-like phages. The short tail appears to have two main gates that retain the genome within the capsid, as previously suggested for the cyanophage P-SCSP1u, for which a structure of the empty particle is unavailable [38]. The Ar-KM primary gate is within the portal protein tunnel, where helix-loops extend inward into the lumen in both the FV and EP, thereby restricting the channel to ∼6 Å. In contrast, the tunnel loop is retracted in EJP, delineating a much larger ∼30 Å lumen that is sufficient to allow ejection proteins to pass first, followed by DNA. Notably, a similar molecular switch to safeguard against genome loss was reported by Bayfield et al. [48], who found that a portal protein structure from the thermophilic phage P23-45 has tunnel loops that adopt a different conformation in the procapsid, thereby acting as a gate. A narrowing of the central lumen was also observed in the crystal structure of phage P22 in a procapsid conformation [49, 50], relative to the mature virion. Therefore, we suggest that the mechanisms of portal protein gating are conserved and rely on the portal functioning like a valve [51]. The similarity between the previously observed procapsid conformation, which occurs before genome packaging, and the post-ejection state suggests that the portal undergoes a complete cycle from a procapsid state to a mature virion [49, 50] and back to a procapsid-like structure after genome ejection. However, the portal is not the only gate that retains the Ar-KM genome within the mature particle. Our study reveals a second gate in the nozzle generated by the protrusion of a short loop. Unlike the portal, which undergoes a local conformational change in the tunnel helix-loop, the secondary gate opens due to a rearrangement of the quaternary structure mediated by gp34 and the ejection protein gp41. In the open tail, the nozzle subunits relax, with β-propellers sliding relative to one another, thereby expanding the central gate. Gp34 inserts at the interface of the nozzle NTI domain, where gp41 forms a helical hairpin. The result is an open nozzle nesting inside a channel formed by gp41 bound to gp42 N-termini. Thus, our work supports the role of gp34 as an ejectosome assembly factor, stabilizing the open nozzle to accommodate gp41 and gp42 after ejection. Previous studies in T7 [18] suggested that gp7.3 serves as a structural scaffold that helps assemble the nozzle into the adapter complex involved in genome ejection, a role that aligns with the proposed function of Ar-KM gp34 as an ejection protein assembly factor. Furthermore, given its putative role as an assembly factor, it is conceivable that the number of gp34 subunits may vary across phages, while retaining a similar underlying function. This may explain the recent finding of 16 copies of gp17, a putative gp7.3-like factor, in the *E. coli* phage E1004 [45]. Or even for phage T7, biochemical measurements of whole T7 virions indicated ∼30 copies of gp7.3 per particle [52]. In contrast, only six copies of gp7.3 were observed within the closed tail of the T7 mature virion [18], but this subunit was not found in ejecting particles with an open tail, similar to Ar-KM EJP described here.

<u>Third</u>, bioinformatic and experimental evidence suggest that Ar-KM is a T4P-specific phage, similar to Luz19 and phiKMV. These phages use the pilus as their initial attachment site; yet, they retain a full complement of ejection proteins reminiscent of those in phage T7, which binds *E. coli* primarily through the lipopolysaccharide core oligosaccharide [15]. Our reconstruction sheds light on the structure of the short tail fiber gp44, while raw micrographs further suggest a network of tentacle-like fibers emanating from the short tail fiber that remain invisible by cryo-EM due to their flexibility. We propose that the flexible Ar-KM tail fiber complex includes additional factors, gp45, gp46, and gp47, all identified in our MS analysis of Ar-KM virions. These putative fibers are likely responsible for pilus attachment and may trigger the conformational changes that initiate deployment of the ejection proteins. We therefore propose two mechanistic models to explain how pilus engagement could be coupled to activation of the ejection machinery during infection. In a first model, the putative fiber complex could wrap around or associate with the T4P in a “jungle-vine” manner, maintaining contact with the pilus while remaining sufficiently mobile to scan the cell envelope. Upon collision with the bacterial surface, a tail-fiber–triggered conformational transition may initiate portal and nozzle opening, leading to the sequential release of the ejection protein assembly factor gp34, followed by gp41, gp42, and ultimately gp43. We further speculate that gp41 may act as a needle-like element that penetrates the OM while remaining associated with gp42, which traverses the periplasm and anchors gp43, the most enigmatic of the ejection proteins. Notably, the peptidoglycan (PG)-hydrolytic activity required to breach the cell wall resides in gp42, Ar-KM’s gp15-like factor, whereas in T7 it is encoded in the N-terminal domain of gp16 [22]. Despite being part of different ejection protein subunits, the PG-degrading lysozyme occupies analogous positions within the assembled DNA-ejectosome (Fig. S5C), underscoring the mosaic evolutionary origins of phage ejection systems.

In an alternative model, attachment of the pilus to the tail fiber complex sends a signal that opens the portal gate and releases the internal proteins gp34 and gp41, similar to how O-antigen binding to phage P22 tailspike triggers tail needle ejection and genome release [25, 53]. Following pilus retraction, interaction between the nozzle and an unknown secondary receptor, most likely deep within the LPS, would facilitate opening of the nozzle gp40, possibly by weakening the numerous intermolecular hydrogen bonds and salt bridges between gp40 subunits, thereby allowing gp34 to wedge itself into place. This, in turn, opens the second gate, releases gp41, which slides out of the nozzle and inserts itself into the OM, and eventually releases gp42 and gp43 to form the PT and IMC required for DNA injection. According to this alternative model, the tail fiber gp44 is not directly involved in recognizing the host surface but instead anchors the pilus, like a fishing hook, to bring the phage to the surface. Future work will clarify the proposed mechanisms of infection.

In summary, this work expands the set of *Pseudomonas* phages solved at near-atomic resolution. It identifies gp34, a gp7.3-like putative ejection-protein assembly factor conserved across diverse phages and essential for T7 viability. Our findings provide mechanistic insight into how phage attachment is coupled to cell-envelope penetration and genome release. The comprehensive 3D atlas of Ar-KM structural proteins established here offers a framework for mapping functional mutations and supports the rational engineering of phages for therapeutic applications.

## MATERIALS AND METHODS

### Origin and characteristics of Ar-KM

Ar-KM was isolated from sewer samples of Southern California. Ar-KM was part of a phage cocktail candidate developed by Armata Pharmaceuticals. Fermentation and purification were performed using proprietary methods in order to achieve clinical levels of purity and a titer of 1 × 10^13^ PFU/ml. The current annotations for Ar-KM were generated from Ar-KM’s published sequence (accession MK837012.1). Glimmer was used to identify additional putative protein-coding sequences and alternative start sites [54]. ARAGORN was used to identify tRNA and tmRNA genes [55]. Sequence alignments of *Bruynogheviruses* from public databases and Armata’s collection were conducted using MAFFT [56] and manually curated to resolve ambiguities with the assumption that ORFs or start codons present in the majority of the phages are more likely to be real than those present in fewer members of the genus. Finally, CDSs were renamed to ensure consistency in their names across members of the phage genus, making future comparisons easier. A list of putative coding sequences is provided in Table S2.

### Vitrification and data collection

2.0 µL of concentrated Ar-KM was applied to a 200-mesh copper Quantifoil R 2/1 holey carbon grid (EMS) that had been glow-discharged for 60 sec at 15 mA using an easiGlow (PELCO). The grid was blotted for 7.5 sec at a blot force of 2 and immediately vitrified in liquid ethane using a Vitrobot Mark IV (Thermo Scientific). Cryo-grids were screened on a 200 kV Glacios (Thermo Scientific) equipped with a Falcon 4 detector (Thermo Scientific) at Thomas Jefferson University. EPU software (Thermo Scientific) was used to collect data in accurate positioning mode. For high-resolution data collection, cryo-micrographs of Ar-KM were acquired on a Titan Krios (Thermo Scientific) microscope operated at 300 kV and equipped with a Falcon 4 direct detector at the Stanford-SLAC CryoEM Center (S2C2).

### Liquid chromatography/mass spectrometry (LC-MS/MS) analysis

Phage samples were treated with 12 mM sodium lauryl sarcosine, 0.5% sodium deoxycholate, and 50 mM triethyl ammonium bicarbonate (TEAB), heated to 95 °C for 10 min, and then sonicated for 10 min, followed by addition 5 mM tris(2-carboxyethyl) phosphine and 10 mM chloroacetamide to reduce, and alkylate the proteins in sample. The sample was then subjected to trypsin digestion overnight (1:100 w/w trypsin added two times). Following digestion, the sample was acidified, lyophilized, and then desalted before injection onto a laser-pulled nanobore C18 column with 1.8 μm beads. This was followed by ionization through a hybrid quadrupole-Orbitrap mass spectrometer. The most abundant proteins were identified by searching the experimental data against a phage protein database, the *Pseudomonas* host protein database, and a common contaminant database using the MASCOT algorithm [57].

### Cryo-EM Single Particle Analysis

*Pseudomonas* phage Ar-KM micrographs were motion-corrected with MotionCorr2 [58]. RELION’s implementation of motion correction was applied to the micrographs using dose-weighted micrographs and the sum of non-dose-weighted power spectra, every 4 e-/Å^2^. Single-particle processing and localized refinements. CTF (Contrast Transfer Function) was estimated in cryoSPARC Patch CTF using default parameters [59]. Owing to the large size of the virions, we initiated manual particle picking (∼200 particles) and generated 2D class templates for subsequent template-based picking. All selected particles were then re-curated by 2D classification. To expedite analysis, particles were split by genome content (DNA-full vs. DNA-empty) and subjected to *ab initio* reconstruction with icosahedral (I) symmetry in parallel. Because the resulting capsids were highly similar, the two sets were merged and refined using non-uniform refinement, and Homogeneous Reconstruction was used to obtain separate full- and empty-capsid maps. For vertex localization, both capsid datasets underwent icosahedral symmetry expansion; a cylindrical mask placed on a C5 axis (generated in ChimeraX [60]) was used for 3D classification to identify tail-containing particles, which were deduplicated, manually re-centered on the tail, re-extracted, and subjected to C6 local refinement. For empty capsids, an additional round of 3D classification of the tail particles separated the tail into two states (EJP and EP). For full capsids, a portal-proximal cylindrical mask revealed unaligned ejection proteins via 3D classification. These particles were re-centered/re-extracted, and refined with C4 local refinement, followed by model-derived tight masking and a second C4 refinement. Blurred density around gp42 and the inner lumen motivated another just-fitting cylindrical mask and non-alignment 3D classification, which explained how the portal barrel breaks symmetry. Unassigned chains were analyzed in ModelAngelo [61], confirming the cross-arranged inner helices as gp34. The tail fibers displayed strong heterogeneity; after C6-symmetry expansion and re-extraction, 3D classification removed particles affected by capsid/DNA overlap (from ∼170,000 to ∼12,000). Knowing the fibers are trimeric with local C3, AlphaFold 3 predictions were used to build masks centered on the cryoSPARC volume alignment; C3 local refinement yielded improved gp44 density. A remaining fuzzy axial helix, consistent with an N-terminal gp45 helix predicted by AlphaFold 3, was isolated with a helical mask for 3D classification to probe the 3 to 1 symmetry conflict; the helix lacked fixed directionality across classes, suggesting high flexibility and orientation-agnostic docking, with the final orientation likely set by an additional conserved contact that is discernible but unassignable at the current resolution. All cryo-EM data-collection statistics are presented in **Table 1**.

### *De novo* model building, oligomer generation, and refinement

All atomic models were built using a combination of ModelAngelo [61], Coot [62], AlphaFold 3 [43], ISOLDE [63], UCSF ChimeraX [64], and Phenix [65]. ModelAngelo was first used to seed well-resolved regions of the high-resolution maps. AlphaFold 3 predictions were segmented in ChimeraX into individual domains, which were then coarsely docked into the density using ChimeraX Matchmaker [64]. These docked fragments were adjusted by rigid-body and real-space fitting in Coot and Phenix, followed by interactive refinement of the carbon backbone and side chains in ISOLDE and Coot.

From the 2.8 Å I-symmetry capsid reconstruction, we built a single icosahedral asymmetric unit (ASU) model. The hexameric major capsid protein was modeled from residues 3–334, whereas the penton subunit could be built for residues 3–221 and 230–334, with the intervening loop remaining unresolved. Four copies of a decoration protein were modeled per ASU; each is present only on hexameric capsomers and could be traced for approximately residues 3–64. Because these decoration copies contact an additional, as-yet unidentified decoration component, they adopt slightly different sequence registers and local conformations, which we modeled as two variants spanning residues 2–58 and 4–59, respectively.

For the FV tail, we likewise built only ASU, but during refinement in Phenix, we imposed C6 symmetry to restrain the fit of side chains and prevent them from drifting into the density of neighboring subunits. Using the 3.2 Å C6-averaged tail map, we modeled the nozzle with a complete sequence except for the C-terminal residue, which lacked interpretable density and was omitted. The HT-adapter was built as a full-length model, tail fiber gp44 was traced from residues 2–251, and the portal was modeled from residues 1–496 for both concentric C6-related rings at the vertex.

We therefore rigid-body fitted the FV tail model into the EP tail map, followed by limited manual adjustments in Coot in a few regions where local differences were apparent. In the EJP, despite the larger conformational change of the nozzle, we applied the same modeling strategy as for the EP nozzle, using the EP model as the starting template. The HT-adapter and tail fibers remain almost unchanged between EP and EJP and were transferred accordingly. The ejection proteins in the EJP state were then built de novo using the same toolbox as for other components, including ModelAngelo [61], AlphaFold 3 [43] with ChimeraX Matchmaker [64], Coot [62], ISOLDE [63], and Phenix [65], yielding models of gp34 (res. 34–81), gp41 (res. 12–153), and gp42 (res. 5–35).

Using the same de novo modeling pipeline, we built the ejection proteins into the 3.5 Å C4-averaged ejection-layer map. In this reconstruction, the C4 gp43 scaffold was modeled as three well-supported segments, spanning residues 74–195, 222–257, and 296–332. Two C4-related copies of gp42 were traced: one from residues 61–881, and a second split into residues 81–603 and 615–881, with the intervening loop remaining unresolved. Likewise, two C4-related gp41 copies were modeled from residues 6–173 and 9–173, respectively. The octameric gp34 was built as eight short helices grouped into two C4-related sets, spanning residues 30–54 and 36–51.

Additionally, a C2-averaged 3.9 Å map was used to further extend and validate the portal barrel. For this map, we abandoned the C6-symmetrized portal model in favor of a C4 description to test whether the symmetry-breaking observed in the ejection-layer map is genuinely encoded in the density. Using the same ejection-protein models as in the C4 ejection reconstruction, we extended three portal subunits beyond the initial 1– 496 model to residues 1–501, 1–505, and 1–510, respectively. These asymmetric extensions of the portal barrel in a C4 framework are consistent with the portal symmetry-breaking mechanism inferred from the localized classifications.

### Structural analysis

All ribbon and surface representations were generated using ChimeraX [64]. Drawings of electron density maps and local resolution maps were generated using ChimeraX [64]. *De novo* density building was attempted using ModelAngelo. Structural neighbors and flexible regions were identified using Foldseek [66]. Binding interfaces were analyzed using PISA [67] to determine bonding interactions, interatomic distances, and bond types. Membrane insertion was predicted using MemBrain [68]. All RMSDs in the Cα position between superimposed structures were calculated using Coot [62].

## Accession numbers

**PDB:** 10YT, 10YU, 10YW, 10ZC, 11AC, 11AE

**EMDB:** 75557, 75558, 75559, 75562, 75582, 75583

## Abbreviations used

cryo-EM: cryogenic electron microscopy
LC-MS/MS: liquid chromatography/mass spectrometry
HT-adapter: head-to-tail adapter
MV: mature virion
EJP: ejecting particle
EP: empty particle
MCP: major capsid protein
PT: periplasmic tunnel
OM: outer membrane
OMC: outer membrane complex
IMC: inner membrane complex
RNAP: RNA polymerase
ASU: asymmetric unit
PG: peptidoglycan
CC: correlation coefficient
RMSD: root-mean-square deviation
dsDNA: double-stranded DNA
SSM: secondary structure superimposition.

## ACKNOWLEDGMENTS

This work was supported by National Institutes of Health grant R35 GM140733 to G.C. Electron microscopy was carried out in the UAB Cryo-EM Facility (RRID:SCR_025450), supported by the Institutional Research Core Program and O’Neal Comprehensive Cancer Center (NIH grant P30 CA013148), with additional funding from NIH grant S10 OD024978. A portion of this work was carried out at Stanford-SLAC CryoEM Center (S2C2), which is supported by the NIH Common Fund Transformative High-Resolution Cryo-Electron Microscopy program (U24 GM129541). The mass spectrometry work was conducted under contract by the Pasarow Mass Spectrometry Laboratory at the Semel Institute for Neuroscience and Human Behavior, University of California, Los Angeles.

## AUTHOR CONTRIBUTIONS STATEMENT

J.L., N.F.B., C-F.D.H., and G.C. performed all steps of the cryo-EM data collection and analysis, deposition of atomic coordinates, and maps. G.C., P.K., D.B., and S.L. supervised the entire project. J.L., N.F.B., and G.C. wrote the paper. N.F.B. and R.K.L. contributed to data analysis, interpretation, and figure creation. S.B., Z.K., R.G., A.S., L.S., and S.L. amplified and purified Ar-KM for cryo-EM analysis. A.S. analyzed the LC-MS/MS data. R.G. and S.L. sequenced and analyzed the genome of Ar-KM. All authors contributed to the writing and editing of the manuscript. All data are available from the corresponding author upon reasonable request.

