## Supplementary Figures and Tables for "Insights into Genome Ejection by a Therapeutic phiKMV-like Bacteriophage"

***Corresponding Author:**

Gino Cingolani, Ph.D.

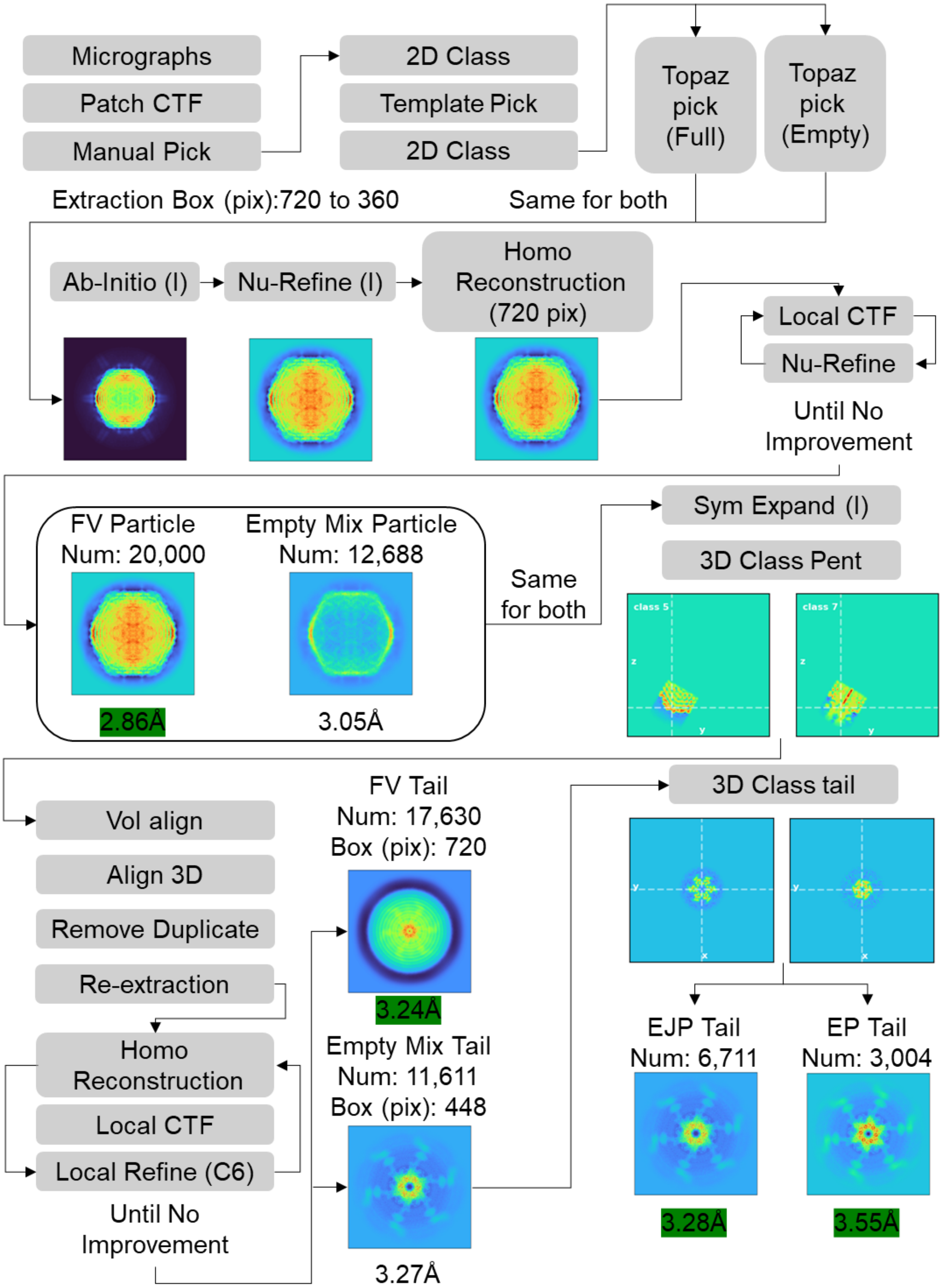


**Figure S1. Flowchart of cryo-EM single-particle analysis used to identify and reconstruct three Ar-KM populations**: full virions (FV), ejecting particles (EJP), and empty particles (EP), all present in the initial dataset of 4,431 movies (a representative micrograph from this dataset is shown in **Fig. 1A**).

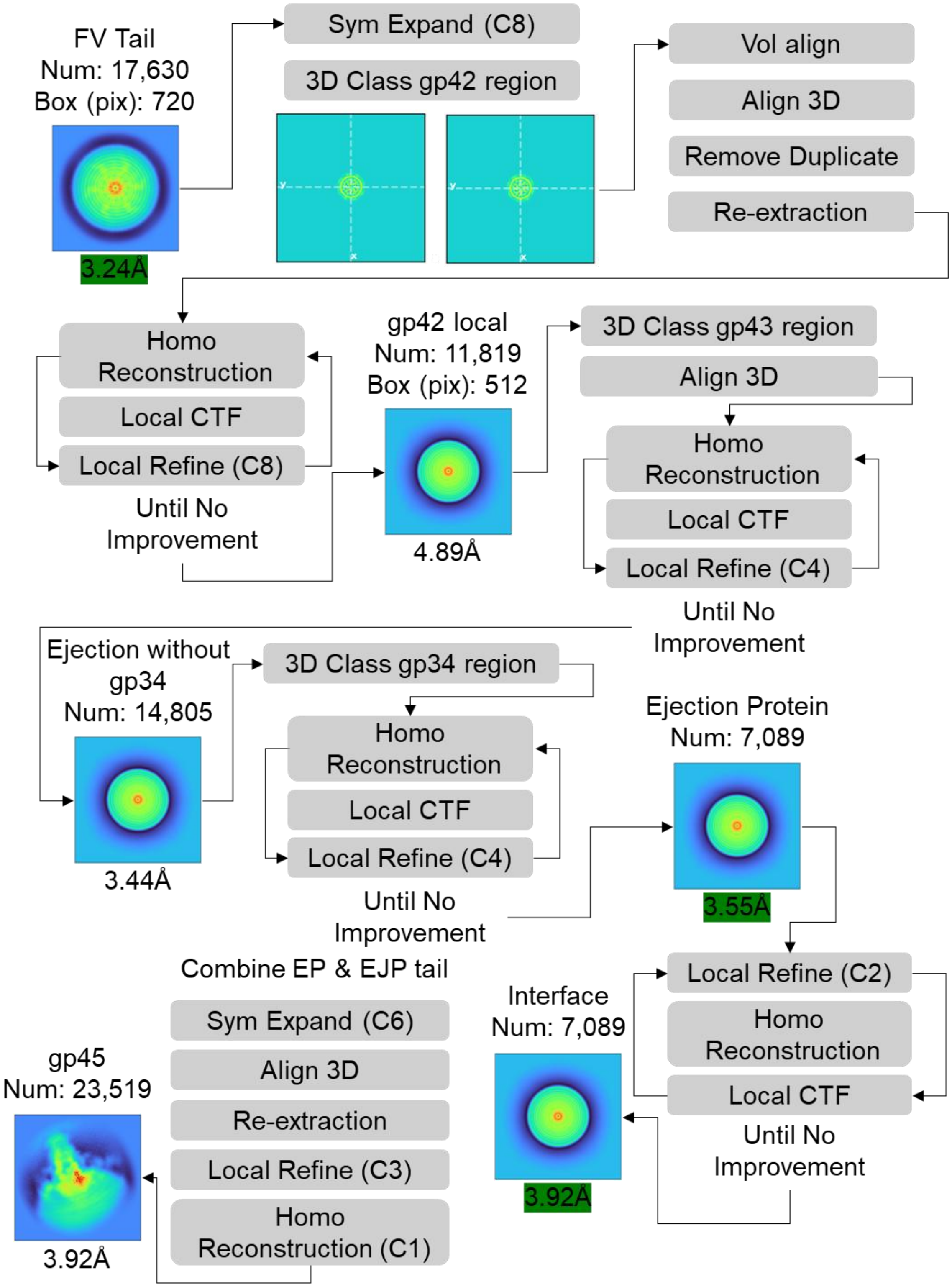


**Figure S2. Flowchart of cryo-EM single-particle analysis used to identify and reconstruct the Ar-KM ejection proteins**. The analysis was conducted using 17,630 FV tail particles (**Table 1**). C2 symmetry was used to visualize the portal barrel:gp41 interface.

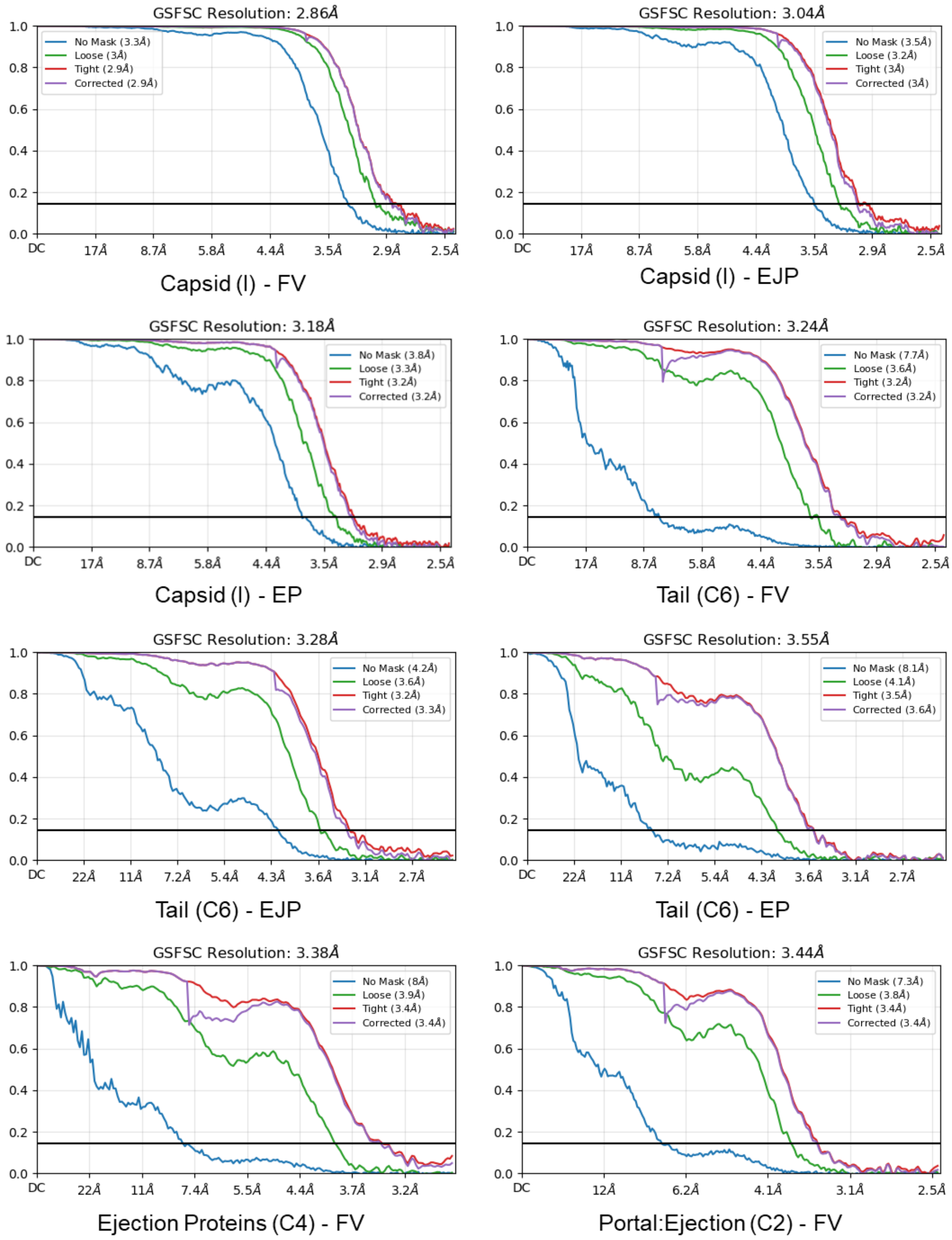


**Figure S3. Fourier Shell Correlation (FSC) resolution curves** for all reconstructions presented in this study. The resolution is indicated at the 0.143 cut-off (generated by cryoSPARC [1]).

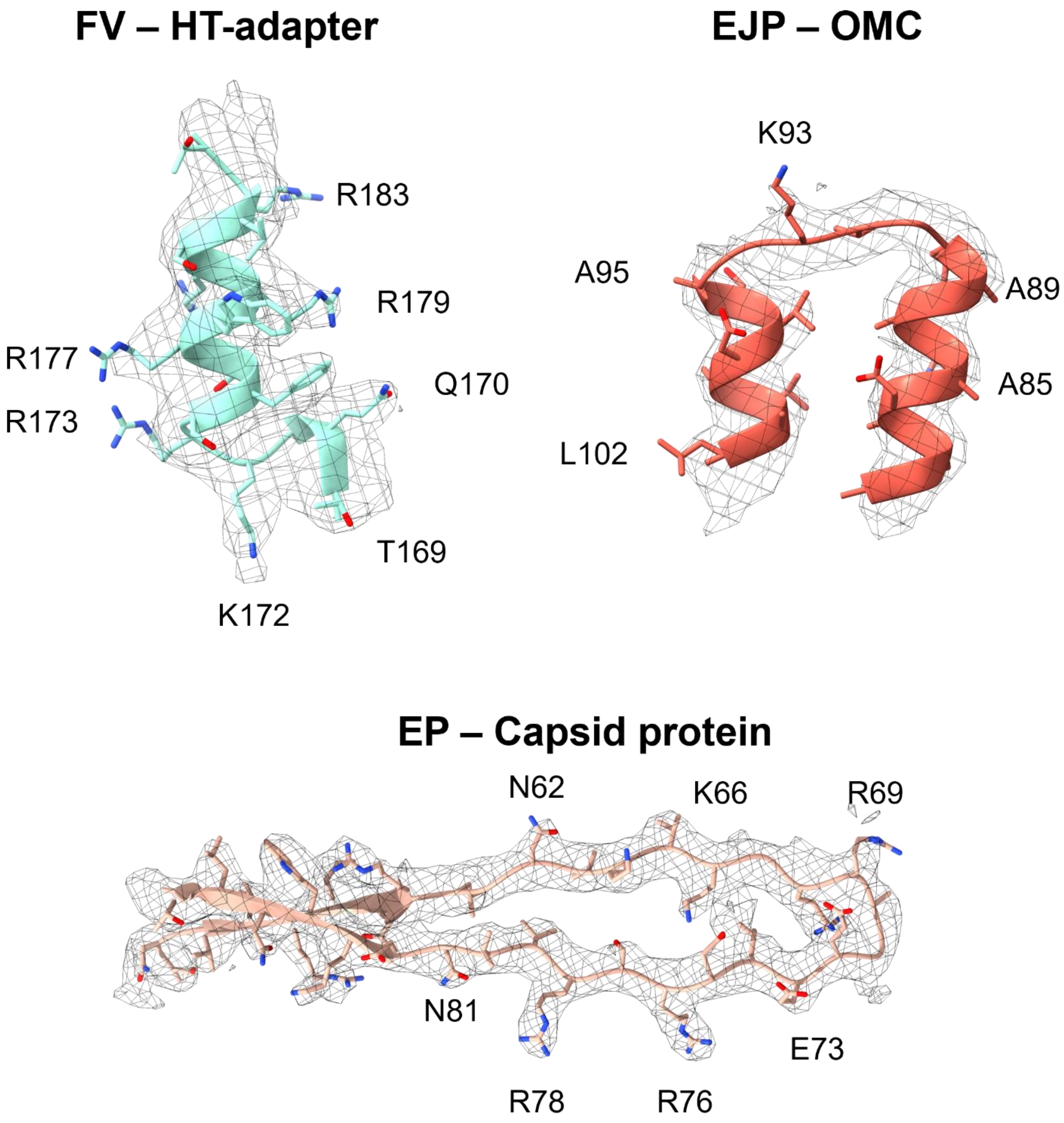


**Figure S4. Representative cryo-EM density.** (Top left) HT-adapter residues 169-184 from FV; (Top right) OMC residues 90-102 from EJP; (bottom) capsid protein residues 55-87 from EP. In all diagrams, the refined atomic model is overlaid on the cryo-EM density, which is displayed at ~3σ contour level. All images were generated using ChimeraX [2].

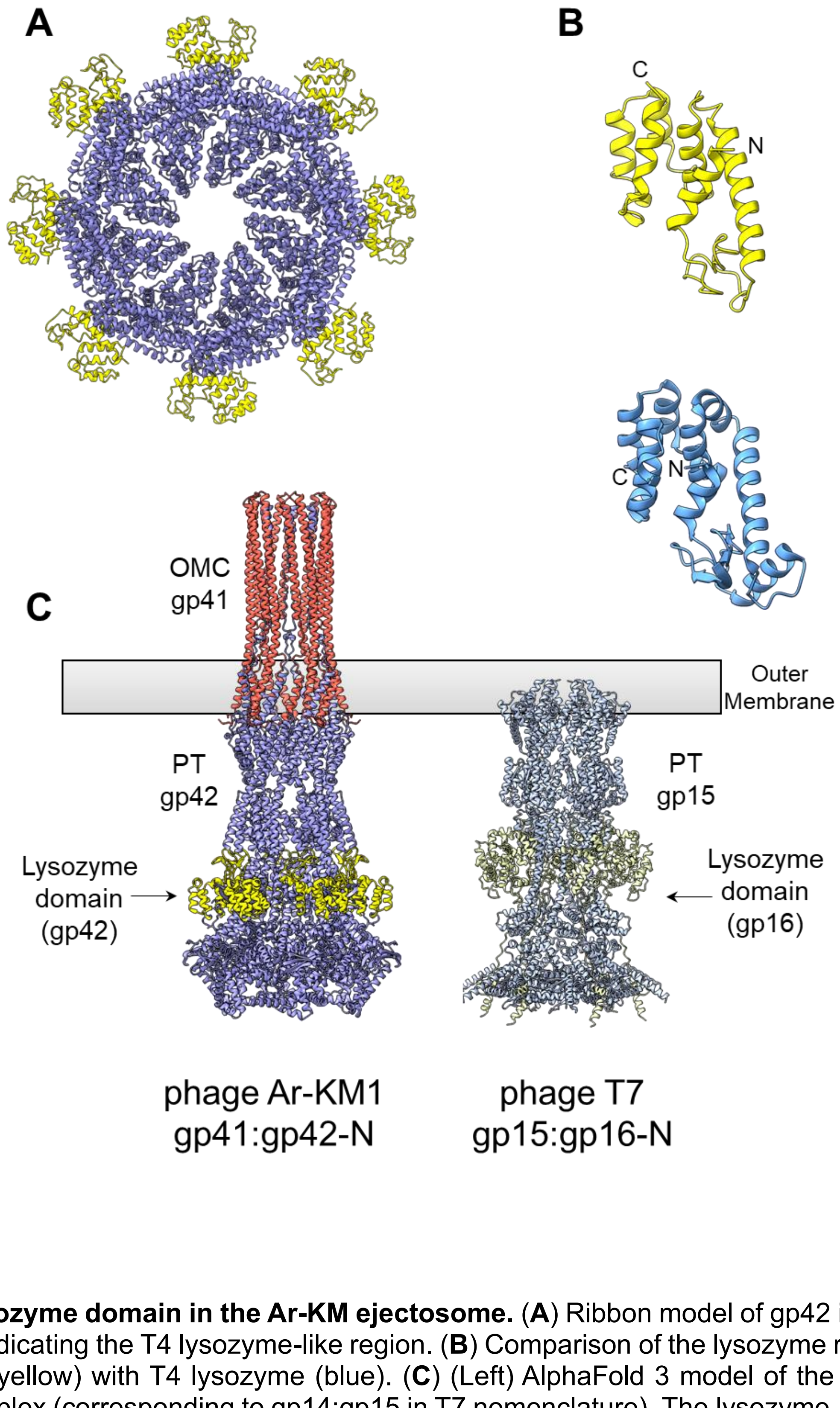


**Figure S5. Lysozyme domain in the Ar-KM ejectosome.** (**A**) Ribbon model of gp42 in FV, with the yellow region indicating the T4 lysozyme-like region. (**B**) Comparison of the lysozyme region of gp42 after trimming (yellow) with T4 lysozyme (blue). (**C**) (Left) AlphaFold 3 model of the phage Ar-KM gp41:gp42 complex (corresponding to gp14:gp15 in T7 nomenclature). The lysozyme-like domain of gp42 is highlighted in yellow. (Right) Phage T7 PT gp15 bound to the N-terminal domain of gp16 (gp16-N) [3], which contains a lysozyme-like domain (yellow).

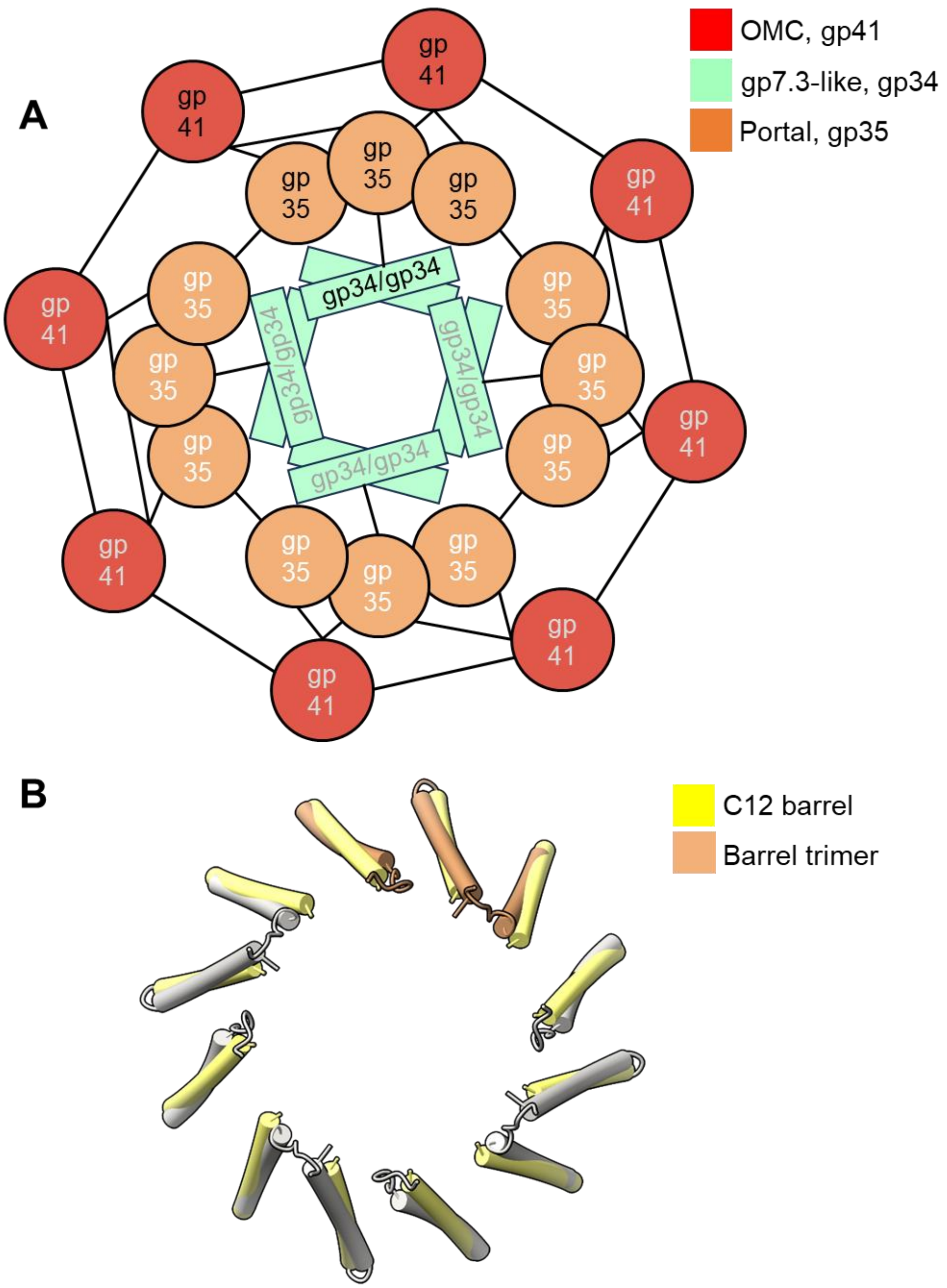


**Figure S6. Schematic of the Ar-KM portal:ejection interface.** (**A**) Schematic diagram of the contacts between the portal barrel (gp35) and the ejection layer formed by gp41 (red) and gp34 (mint). (**B**) Comparison between the theoretical position of the C12 portal barrel helices (egg-yolk yellow) and the observed barrel helices (gray and tan), revealing significant deviations from C12 symmetry. Only three barrel helices from the cryo-EM reconstruction are colored egg-yolk yellow, while the other nine barrel helices are gray.

### Table S1. Conservation of Ar-KM proteins identified in this study across 60 phiKMV-like *viruses*

| Phage name | Accession number | Overall phage genome identity to Ar-KM | Protein identity to Ar-KM | | | | | | | | | | | | | | | |
|---|---|---|---|---|---|---|---|---|---|---|---|---|---|---|---|---|---|---|
| | | | gp34 | gp35 | gp36 | gp37 | gp39 | gp40 | gp41 | gp42 | gp43 | gp44 | gp45 | gp46 | | gp47 | gp54 | gp55 |
| | | | | | | | | | | | | | | N-Ter 1-175 | C-Ter 154-302 | | | |
| 130_113 | MH107770 | 92.2 | 100.0 | 99.8 | 99.7 | 97.0 | 99.5 | 99.5 | 98.9 | 99.7 | 99.3 | 99.6 | 94.7 | 98.0 | | 99.5 | 96.2 | 88.7 |
| vB_Pae_QDWS | MZ687409 | 91.4 | 100.0 | 99.2 | 98.1 | 97.0 | 99.5 | 98.4 | 97.8 | 99.4 | 99.1 | 97.6 | 65.8 | 74.5 | | 88.6 | 98.1 | 86.6 |
| vB_Pae4841-AFR43 | PP501790 | 91.4 | 98.0 | 99.6 | 97.8 | 97.0 | 100.0 | 97.6 | 99.4 | 99.1 | 99.1 | 98.8 | 94.7 | 98.3 | | 88.6 | 99.0 | 89.2 |
| HX1 | MW406976 | 91.3 | 100.0 | 99.6 | 97.8 | 97.0 | 100.0 | 98.3 | 96.7 | 99.4 | 99.2 | 98.0 | 90.8 | 98.3 | | 88.6 | 98.1 | 90.8 |
| RLP | MH979674 | 91.3 | 100.0 | 99.8 | 96.0 | 97.0 | 98.9 | 98.3 | 98.9 | 99.2 | 99.3 | 98.0 | 94.7 | 97.7 | | 88.1 | 99.0 | 89.7 |
| Ps23.FT | PV641621 | 91 | 100.0 | 99.4 | 98.1 | 97.0 | 99.5 | 98.3 | 98.9 | 99.1 | 99.2 | 97.6 | 95.4 | 98.0 | | 88.6 | 98.1 | 89.5 |
| 20176-10 | PQ510339 | 90.9 | 100.0 | 99.4 | 97.8 | 97.0 | 100.0 | 98.2 | 97.8 | 99.3 | 99.3 | 98.8 | 94.7 | 98.3 | | 88.6 | 99.0 | 89.2 |
| 20Aug401 | OQ319930 | 90.9 | 99.0 | 99.4 | 97.5 | 97.0 | 98.4 | 98.8 | 99.4 | 99.2 | 98.7 | 98.0 | 90.8 | 97.7 | | 88.1 | 97.1 | 86.6 |
| PAO1-15pyo | LN610580 | 90.9 | 98.0 | 99.2 | 97.5 | 97.0 | 97.3 | 98.6 | 99.4 | 99.2 | 98.7 | 94.8 | 90.8 | 96.0 | | 97.5 | 94.2 | 87.2 |
| vB_PaeA_SB | OR208619 | 90.9 | 100.0 | 99.6 | 97.5 | 98.5 | 99.5 | 98.1 | 99.4 | 99.1 | 99.0 | 98.0 | 94.7 | 99.7 | | 98.5 | 98.1 | 91.8 |
| vB_PaeM_FBPa3 | ON857926 | 90.9 | 100.0 | 99.0 | 96.6 | 94.6 | 100.0 | 98.3 | 98.9 | 99.2 | 99.1 | 98.0 | 52.0 | 70.2 | | 99.0 | 98.1 | 87.5 |
| CP-p-PA-21037 | PV844377 | 90.8 | 98.0 | 99.0 | 97.8 | 97.0 | 97.3 | 98.8 | 99.4 | 99.2 | 98.6 | 94.4 | 90.8 | 96.0 | | 98.0 | 98.1 | 85.1 |
| 20176-1 | PQ510338 | 90.7 | 98.0 | 99.6 | 97.8 | 97.0 | 100.0 | 97.6 | 99.4 | 99.1 | 99.1 | 98.4 | 90.8 | 98.7 | | 88.6 | 99.0 | 89.2 |
| PAXYB1 | KY618819 | 90.7 | 100.0 | 99.2 | 97.8 | 97.0 | 100.0 | 98.3 | 99.4 | 99.3 | 99.1 | 98.0 | 52.0 | 70.5 | | 99.0 | 98.1 | 90.6 |
| vB_Pae_PLY | OR689712 | 90.6 | 100.0 | 99.4 | 96.6 | 97.0 | 98.4 | 98.9 | 98.9 | 99.1 | 99.2 | 98.0 | 94.7 | 98.3 | | 88.6 | 98.1 | 87.4 |
| Phipa2 | OK539824 | 90.4 | 99.0 | 99.2 | 96.9 | 96.7 | 98.4 | 98.4 | 99.4 | 99.2 | 99.2 | 98.0 | 95.4 | 99.0 | | 88.6 | 97.1 | 86.8 |
| vB_Pae10145-KEN1 | PP501791 | 90.3 | 99.0 | 99.4 | 97.8 | 97.3 | 100.0 | 98.8 | 99.4 | 99.3 | 99.4 | 98.4 | 90.8 | 98.3 | | 88.6 | 98.1 | 87.7 |
| L5 | OL754589 | 90.2 | 100.0 | 99.2 | 97.8 | 96.4 | 99.5 | 98.3 | 97.2 | 99.2 | 99.2 | 98.0 | 65.8 | 73.8 | | 88.6 | 98.1 | 87.8 |
| MPK6 | NC_022746 | 90.2 | 100.0 | 99.4 | 97.8 | 97.0 | 99.5 | 98.3 | 98.9 | 98.4 | 99.0 | 98.0 | 52.6 | 70.5 | | 99.0 | 98.1 | 82.0 |
| DL62 | NC_028836 | 90.1 | 99.0 | 99.4 | 97.2 | 97.0 | 98.9 | 98.9 | 98.3 | 99.6 | 98.9 | 97.6 | 90.1 | 99.3 | | 88.6 | 73.1 | 72.0 |
| CP-p-PA-22060 | PQ859843 | 90 | 100.0 | 99.8 | 97.8 | 94.9 | 100.0 | 98.2 | 97.8 | 99.4 | 99.1 | 97.6 | 90.1 | 98.7 | | 88.6 | 98.1 | 84.6 |
| tbilissiM32 | NC_017865 | 89.9 | 100.0 | 99.6 | 97.8 | 97.0 | 97.3 | 98.7 | 99.4 | 99.2 | 98.6 | 94.8 | 90.8 | 96.0 | | 98.0 | 95.2 | 87.0 |
| vB_Pae9718-KEN10 | PP501792 | 89.6 | 100.0 | 99.8 | 97.5 | 97.0 | 100.0 | 98.3 | 98.9 | 99.2 | 99.1 | 98.0 | 90.8 | 98.7 | | 88.6 | 98.1 | 86.7 |
| 20176-2 | PQ510340 | 89.5 | 100.0 | 99.4 | 97.8 | 97.0 | 100.0 | 97.8 | 99.4 | 99.1 | 99.1 | 98.8 | 94.7 | 98.3 | | 88.6 | 99.0 | 89.2 |
| PT5 | EU056923 | 89.5 | 98.0 | 99.2 | 97.8 | 97.0 | 97.8 | 98.5 | 99.4 | 99.2 | 98.7 | 94.8 | 90.8 | 96.4 | | 97.5 | 95.2 | 86.7 |
| JB10 | OQ412633 | 89.3 | 99.0 | 100.0 | 97.8 | 98.8 | 97.3 | 98.4 | 98.9 | 99.7 | 99.4 | 94.8 | 26.0 | 24.8 | | 47.5 | 98.1 | 49.6 |
| vB_PaeP_PE3 | MN901924 | 89.2 | 99.0 | 99.4 | 96.9 | 97.0 | 98.4 | 99.0 | 99.4 | 99.0 | 99.0 | 98.0 | 52.6 | 69.9 | | 99.0 | 39.5 | 50.7 |
| ASP23 | MN602045 | 89.1 | 98.0 | 99.2 | 97.8 | 97.0 | 97.3 | 98.8 | 99.4 | 99.2 | 98.7 | 95.6 | 26.0 | 24.5 | | 50.5 | 95.2 | 87.0 |
| LUZ19 | NC_010326 | 89.1 | 100.0 | 99.4 | 97.8 | 97.0 | 98.4 | 99.0 | 99.4 | 99.1 | 99.1 | 98.0 | 52.6 | 69.5 | | 99.0 | 97.1 | 51.7 |
| vB_PaeP_SPCB | MN615698 | 89.1 | 96.9 | 99.2 | 97.8 | 97.0 | 97.3 | 98.4 | 99.4 | 99.4 | 98.6 | 94.4 | 90.8 | 96.4 | | 97.5 | 95.2 | 86.7 |
| vB_PaeA_55_1W | MZ553931 | 88.9 | 100.0 | 99.4 | 97.8 | 97.0 | 98.4 | 99.0 | 99.4 | 99.3 | 98.9 | 98.0 | 52.0 | 70.2 | | 98.5 | 96.2 | 96.9 |
| MYY9 | MW406975 | 88.8 | 100.0 | 99.2 | 96.6 | 97.0 | 100.0 | 98.4 | 98.9 | 99.4 | 99.3 | 98.0 | 52.6 | 70.5 | | 99.0 | 97.1 | 88.2 |
| vB_PaBD_211 | PV786137 | 88.7 | 100.0 | 99.4 | 98.1 | 97.0 | 100.0 | 98.4 | 99.4 | 99.4 | 99.4 | 95.6 | 27.3 | 24.5 | | 47.5 | 97.1 | 84.8 |
| vB_PaPhi_Mx1 | OR594342 | 88.7 | 99.0 | 99.2 | 98.8 | 97.0 | 98.9 | 98.1 | 97.2 | 99.4 | 99.3 | 98.0 | 52.6 | 51.7 | 72.5 | 98.5 | 96.2 | 52.9 |
| vB_PaeP_SPCG | MN615699 | 88.6 | 96.9 | 99.2 | 97.8 | 97.0 | 97.3 | 98.4 | 99.4 | 99.2 | 98.7 | 94.4 | 90.8 | 96.7 | | 98.0 | 95.2 | 86.3 |
| vB_PaePA8132phi1_PS3 | PV565006 | 88.6 | 100.0 | 99.8 | 97.8 | 97.0 | 100.0 | 98.8 | 98.9 | 99.7 | 99.3 | 98.4 | 90.8 | 98.3 | | 88.6 | 98.1 | 89.3 |
| vB PaM SEMA | OP556577 | 88.5 | 100.0 | 99.4 | 98.1 | 94.9 | 97.8 | 98.7 | 98.9 | 99.4 | 99.2 | 97.6 | 52.6 | 70.2 | | 98.5 | 94.2 | 74.5 |
| MBL | KC969441 | 88.4 | 100.0 | 99.4 | 97.2 | 97.0 | 97.3 | 98.8 | 99.4 | 99.2 | 98.7 | 95.6 | 26.0 | 24.5 | | 50.5 | 98.1 | 83.9 |
| PaFZ4 | PV610699 | 88.3 | 100.0 | 100.0 | 97.5 | 96.7 | 99.5 | 98.3 | 97.8 | 99.3 | 99.3 | 94.8 | 26.7 | 22.3 | | 47.5 | 97.1 | 86.6 |
| Henu_18-20N | PV864296 | 88.2 | 100.0 | 99.6 | 97.5 | 96.7 | 100.0 | 98.3 | 99.4 | 99.3 | 99.2 | 95.6 | 27.3 | 24.5 | | 47.5 | 98.1 | 74.7 |
| MPK7 | NC_022091 | 88.2 | 100.0 | 99.4 | 97.5 | 97.0 | 97.8 | 98.8 | 97.8 | 99.3 | 99.0 | 95.6 | 27.3 | 24.5 | | 47.5 | 98.1 | 87.2 |
| Ab05 | NC_026602 | 88.1 | 100.0 | 99.4 | 97.8 | 97.0 | 98.9 | 98.3 | 97.2 | 94.1 | 98.3 | 97.6 | 90.8 | 97.4 | | 88.6 | 74.0 | 82.7 |
| PT2 | NC_011107 | 88.1 | 100.0 | 99.4 | 97.2 | 97.0 | 97.3 | 98.8 | 99.4 | 99.2 | 98.7 | 95.6 | 26.0 | 24.5 | | 50.5 | 98.1 | 84.1 |
| vB_PaeP_FBPa25 | ON857935 | 87.4 | 100.0 | 99.0 | 96.6 | 94.6 | 100.0 | 98.3 | 98.9 | 96.9 | 90.7 | 97.3 | 52.0 | 70.2 | | 99.0 | 98.1 | 87.6 |
| vB_PaeP_PPA-ABTNL | NC_027375 | 87.4 | 100.0 | 99.2 | 95.3 | 97.0 | 100.0 | 98.8 | 98.3 | 99.3 | 99.1 | 95.6 | 27.3 | 24.5 | | 47.5 | 98.1 | 86.8 |
| BARC01 | PQ641084 | 87.3 | 100.0 | 97.1 | 92.0 | 95.2 | 98.4 | 97.7 | 91.2 | 98.1 | 99.0 | 98.8 | 90.1 | 98.3 | | 88.6 | 72.1 | 84.8 |
| phiLP | LC727700 | 87.2 | 100.0 | 99.2 | 98.1 | 97.0 | 98.4 | 98.2 | 97.8 | 99.4 | 99.0 | 95.6 | 27.3 | 24.2 | | 47.5 | 98.1 | 88.1 |
| phiNFS | KU743887 | 87.1 | 98.0 | 99.2 | 97.8 | 97.0 | 97.3 | 98.3 | 99.4 | 99.3 | 98.7 | 95.6 | 26.0 | 24.5 | | 50.5 | 95.2 | 87.0 |
| vB_PaeP_P1G | OQ230793 | 87.1 | 99.0 | 99.4 | 98.4 | 97.0 | 98.4 | 99.2 | 98.3 | 95.2 | 98.7 | 98.0 | 52.6 | 70.5 | | 98.5 | 97.1 | 88.6 |
| vB_PaePA10145phi1_HR2 | PV565007 | 87.1 | 99.0 | 100.0 | 97.5 | 97.0 | 98.9 | 99.2 | 98.9 | 99.3 | 99.2 | 98.0 | 90.8 | 98.3 | | 88.6 | 72.1 | 76.0 |
| PA69 | OR238910 | 87 | 100.0 | 99.6 | 97.8 | 97.0 | 99.5 | 98.3 | 97.8 | 99.3 | 99.3 | 98.0 | 94.1 | 97.4 | | 88.6 | 97.1 | 87.1 |
| phiKMV | NC_005045 | 87 | 98.0 | 99.2 | 97.8 | 97.0 | 97.3 | 98.8 | 99.4 | 99.1 | 98.7 | 95.6 | 26.0 | 24.5 | | 50.5 | 96.2 | 87.0 |
| phikF77 | NC_012418 | 86.8 | 98.0 | 98.2 | 92.0 | 94.9 | 97.8 | 97.5 | 97.8 | 98.0 | 97.8 | 98.0 | 52.6 | 69.5 | | 99.0 | 96.2 | 89.1 |
| vB_PaeP_FBPa18 | ON857933 | 86.2 | 100.0 | 99.0 | 96.6 | 94.6 | 100.0 | 98.3 | 98.9 | 99.2 | 99.1 | 98.0 | 52.0 | 52.9 | | 99.0 | 98.1 | 88.0 |
| PNM | OP292288 | 86.1 | 98.0 | 99.2 | 97.8 | 97.0 | 98.4 | 98.1 | 99.4 | 99.1 | 98.4 | 95.2 | 26.0 | 24.8 | | 50.5 | 98.1 | 85.1 |
| vB_PaeP_FBPa6 | ON857928 | 86 | 100.0 | 99.0 | 96.6 | 94.6 | 100.0 | 98.3 | 98.9 | 99.2 | 99.1 | 98.0 | 52.0 | 58.3 | | 99.0 | 98.1 | 87.6 |
| AIIMS-Pa-A1 | MW117144 | 84.4 | 100.0 | 99.4 | 97.5 | 97.0 | 100.0 | 95.2 | 93.4 | 93.3 | 98.0 | 94.0 | 89.5 | 98.3 | | 88.6 | 94.2 | 95.1 |
| vB_PaePA01phi1_RS1 | PV565004 | 83.7 | 100.0 | 99.4 | 97.8 | 97.0 | 100.0 | 98.5 | 99.4 | 99.4 | 99.5 | 99.6 | 90.8 | 98.0 | | 88.6 | 97.1 | 85.3 |
| LKD16 | NC_009935 | 82.8 | 98.0 | 96.9 | 90.5 | 96.4 | 98.4 | 94.7 | 92.3 | 99.1 | 98.8 | 99.6 | 65.8 | 75.5 | | 88.1 | 99.0 | 86.6 |
| PJNP013 | OR941786 | 78.8 | 98.0 | 96.9 | 91.7 | 94.9 | 98.4 | 94.4 | 92.3 | 97.2 | 97.8 | 99.2 | 63.2 | 75.2 | | 88.1 | 74.0 | 70.3 |

**Table S2. Conservation of ejection proteins across 205 genomes of phages in the order *Autographivirales*.**

| | OMC | | PT | | | | IMC | | | |
|---|---|---|---|---|---|---|---|---|---|---|
| **Accession #** | **OMC length (AA)** | **OMC domain** | **PT Length (AA)** | **PT domain** | **location and type of cell wall degradation domain** | | | | **IMC length (AA)** | **IMC domain** |
| | | | | | **middle** | **C-term** | **N-term** | **middle** | | |
| OR915848.1 | 198 | IPR038996 | 751 | IPR038993 | N/A | N/A | IPR000189 IPR023346 | N/A | 1323 | IPR038994 |
| OR822025.1 | 205 | IPR038996 | 754 | IPR038993 | N/A | N/A | IPR000189 IPR023346 | N/A | 1326 | IPR038994 |
| KY981272.1 | 242 | N/A | 783 | N/A | N/A | N/A | IPR002901 | N/A | 1578 | N/A |
| AB920995.1 | 326 | N/A | 759 | N/A | N/A | N/A | IPR008258 | N/A | 1373 | N/A |
| MW960043.1 | 277 | N/A | 815 | N/A | N/A | N/A | IPR008258 | N/A | 1361 | N/A |
| MW980070.1 | 179 | IPR038996 | 675 | N/A | N/A | N/A | IPR008258 IPR023346 | N/A | 1259 | N/A |
| JX483873.1 | 181 | IPR038996 | 675 | N/A | N/A | N/A | IPR008258 IPR023346 | N/A | 1258 | N/A |
| PP226939.1 | 193 | IPR038996 | 686 | IPR038993 | N/A | N/A | IPR008258 IPR023346 | N/A | 1361 | N/A |
| ON604651.1 | 198 | IPR038996 | 687 | IPR038993 | N/A | N/A | IPR008258 IPR023346 | N/A | 1354 | N/A |
| OW991346.2 | 190 | IPR038996 | 693 | IPR038993 | N/A | N/A | IPR008258 IPR023346 | N/A | 1431 | N/A |
| MT740748.1 | 187 | IPR038996 | 698 | IPR038993 | N/A | N/A | IPR008258 IPR023346 | N/A | 1335 | N/A |
| KY626176.1 | 182 | IPR038996 | 710 | N/A | N/A | N/A | IPR008258 IPR023346 | N/A | 1402 | N/A |
| MH179472.2 | 181 | IPR038996 | 716 | IPR038993 | N/A | N/A | IPR008258 IPR023346 | N/A | 1334 | N/A |

| | | | | | | | | | | |
|---|---|---|---|---|---|---|---|---|---|---|
| MF893341.1 | 175 | IPR038996 | 723 | IPR038993 | N/A | N/A | IPR008258 IPR023346 | N/A | 1296 | N/A |
| KC960671.1 | 198 | IPR038996 | 724 | IPR038993 | N/A | N/A | IPR008258 IPR023346 | N/A | 1319 | IPR038994 |
| KT949345.1 | 265 | N/A | 725 | N/A | N/A | N/A | IPR008258 IPR023346 | N/A | 1253 | N/A |
| KR153873.1 | 191 | IPR038996 | 727 | IPR038993 | N/A | N/A | IPR008258 IPR023346 | N/A | 1291 | N/A |
| AB597179.1 | 191 | IPR038996 | 729 | IPR038993 | N/A | N/A | IPR008258 IPR023346 | N/A | 1291 | N/A |
| KP343639.1 | 195 | IPR038996 | 729 | IPR038993 | N/A | N/A | IPR008258 IPR023346 | N/A | 1295 | N/A |
| MG775258.1 | 194 | IPR038996 | 731 | IPR038993 | N/A | N/A | IPR008258 IPR023346 | N/A | 1400 | N/A |
| OQ716796.1 | 196 | N/A | 732 | IPR038993 | N/A | N/A | IPR008258 IPR023346 | N/A | 1335 | N/A |
| KX431888.1 | 186 | IPR038996 | 733 | IPR038993 | N/A | N/A | IPR008258 IPR023346 | N/A | 1331 | N/A |
| MT711889.1 | 190 | IPR038996 | 733 | IPR038993 | N/A | N/A | IPR008258 IPR023346 | N/A | 1394 | N/A |
| OM131411.1 | 192 | IPR038996 | 733 | IPR038993 | N/A | N/A | IPR008258 IPR023346 | N/A | 1338 | N/A |
| FR823298.1 | 183 | IPR038996 | 734 | IPR038993 | N/A | N/A | IPR008258 IPR023346 | N/A | 1327 | N/A |
| OQ622254.1 | 189 | IPR038996 | 735 | IPR038993 | N/A | N/A | IPR008258 IPR023346 | N/A | 1350 | N/A |
| OQ622093.1 | 187 | IPR038996 | 737 | IPR038993 | N/A | N/A | IPR008258 IPR023346 | N/A | 1398 | N/A |

| | | | | | | | | | | |
|---|---|---|---|---|---|---|---|---|---|---|
| KP025626.1 | 186 | IPR038996 | 738 | IPR038993 | N/A | N/A | IPR008258 IPR023346 | N/A | 1332 | N/A |
| MF166859.1 | 179 | IPR038996 | 739 | IPR038993 | N/A | N/A | IPR008258 IPR023346 | N/A | 1362 | IPR038994 |
| AF493143.1 | 194 | IPR038996 | 739 | IPR038993 | N/A | N/A | IPR008258 IPR023346 | N/A | 1393 | N/A |
| KX756572.1 | 256 | N/A | 744 | N/A | N/A | N/A | IPR008258 IPR023346 | N/A | 1269 | N/A |
| JN651747.1 | 260 | N/A | 745 | N/A | N/A | N/A | IPR008258 IPR023346 | N/A | 1250 | N/A |
| KT184661.1 | 258 | N/A | 746 | N/A | N/A | N/A | IPR008258 IPR023346 | N/A | split 784+480 | N/A |
| KC853746.1 | 286 | N/A | 747 | N/A | N/A | N/A | IPR008258 IPR023346 | N/A | 1346 | N/A |
| V01146.1 | 197 | IPR038996 | 748 | IPR038993 | N/A | N/A | IPR008258 IPR023346 | N/A | 1319 | IPR038994 |
| MT682386.1 | 252 | IPR038996 | 749 | N/A | N/A | N/A | IPR008258 IPR023346 | N/A | 1255 | N/A |
| MN509793.1 | 255 | N/A | 749 | N/A | N/A | N/A | IPR008258 IPR023346 | N/A | 1261 | N/A |
| HE956710.2 | 256 | IPR038996 | 750 | N/A | N/A | N/A | IPR008258 IPR023346 | N/A | 1260 | N/A |
| MT597419.1 | 183 | IPR038996 | 751 | IPR038993 | N/A | N/A | IPR008258 IPR023346 | N/A | 1372 | N/A |
| LT960606.1 | 200 | IPR038996 | 751 | IPR038993 | N/A | N/A | IPR008258 IPR023346 | N/A | 1318 | N/A |

| | | | | | | | | | | |
|---|---|---|---|---|---|---|---|---|---|---|
| LC779549.1 | 264 | IPR038996 | 751 | N/A | N/A | N/A | IPR008258 IPR023346 | N/A | 1266 | N/A |
| PP453689.1 | 264 | N/A | 751 | N/A | N/A | N/A | IPR008258 IPR023346 | N/A | 1263 | N/A |
| MF621978.1 | 411 | N/A | 752 | N/A | N/A | N/A | IPR008258 IPR023346 | N/A | 1318 | N/A |
| HE956707.2 | 266 | N/A | 753 | N/A | N/A | N/A | IPR008258 IPR023346 | N/A | 1254 | N/A |
| KY250035.2 | 194 | IPR038996 | 754 | IPR038993 | N/A | N/A | IPR008258 IPR023346 | N/A | 1339 | N/A |
| MN013082.1 | 198 | IPR038996 | 755 | IPR038993 | N/A | N/A | IPR008258 IPR023346 | N/A | 1325 | N/A |
| MH059637.1 | 205 | IPR038996 | 755 | IPR038993 | N/A | N/A | IPR008258 IPR023346 | N/A | 1328 | IPR038994 |
| MN184885.1 | 194 | IPR038996 | 757 | IPR038993 | N/A | N/A | IPR008258 IPR023346 | N/A | 1316 | N/A |
| MF979561.1 | 208 | IPR038996 | 758 | IPR038993 | N/A | N/A | IPR008258 IPR023346 | N/A | 1329 | IPR038994 |
| MK290739.2 | 208 | IPR038996 | 760 | IPR038993 | N/A | N/A | IPR008258 IPR023346 | N/A | 1332 | IPR038994 |
| MZ326863.1 | 304 | N/A | 762 | N/A | N/A | N/A | IPR008258 IPR023346 | N/A | 1641 | N/A |
| JN882298.1 | 276 | IPR038996 | 770 | N/A | N/A | N/A | IPR008258 IPR023346 | N/A | 1401 | N/A |
| KF626666.1 | 284 | N/A | 774 | N/A | N/A | N/A | IPR008258 IPR023346 | N/A | 1364 | N/A |

| | | | | | | | | | | |
|---|---|---|---|---|---|---|---|---|---|---|
| MW965453.2 | 337 | N/A | 774 | N/A | N/A | N/A | IPR008258 IPR023346 | N/A | 1602 | N/A |
| LR778216.1 | 299 | N/A | 776 | N/A | N/A | N/A | IPR008258 IPR023346 | N/A | 1491 | N/A |
| MH684921.1 | 251 | IPR038996 | 780 | N/A | N/A | N/A | IPR008258 IPR023346 | N/A | 1300 | N/A |
| MF979559.1 | 173 | IPR038996 | 781 | | N/A | N/A | IPR008258 IPR023346 | N/A | 1574 | N/A |
| KX660669.1 | 273 | N/A | 781 | N/A | N/A | N/A | IPR008258 IPR023346 | N/A | 1320 | N/A |
| MH059632.2 | 195 | IPR038996 | 782 | IPR038993 | N/A | N/A | IPR008258 IPR023346 | N/A | 1321 | N/A |
| AB451219.1 | 287 | N/A | 786 | N/A | N/A | N/A | IPR008258 IPR023346 | N/A | 1607 | N/A |
| KF626667.1 | 295 | N/A | 787 | N/A | N/A | N/A | IPR008258 IPR023346 | N/A | 1633 | N/A |
| KY316062.1 | 307 | IPR038996 | 788 | N/A | N/A | N/A | IPR008258 IPR023346 | N/A | 1690 | N/A |
| MT740728.1 | 307 | N/A | 794 | N/A | N/A | N/A | IPR008258 IPR023346 | N/A | 1589 | N/A |
| MH370477.2 | 175 | IPR038996 | 806 | IPR038993 | N/A | N/A | IPR008258 IPR023346 | N/A | 1321 | N/A |
| GU393987.1 | 269 | N/A | 809 | N/A | N/A | N/A | IPR008258 IPR023346 | N/A | 1334 | N/A |
| PP079415.1 | 275 | N/A | 812 | N/A | N/A | N/A | IPR008258 IPR023346 | N/A | 1354 | N/A |
| MK903280.1 | 283 | N/A | 821 | N/A | N/A | N/A | IPR008258 IPR023346 | N/A | 1361 | N/A |

| | | | | | | | | | | |
|---|---|---|---|---|---|---|---|---|---|---|
| KT321314.1 | 183 | IPR038996 | 747 | IPR038993 | N/A | N/A | IPR008258 IPR023346 IPR000189 | N/A | 1316 | N/A |
| EU734173.1 | 197 | IPR038996 | 752 | IPR038993 | N/A | N/A | IPR008258 IPR023346 IPR000189 | N/A | 1322 | N/A |
| MG966531.1 | 197 | IPR038996 | 752 | IPR038993 | N/A | N/A | IPR008258 IPR023346 IPR000189 | N/A | 1322 | N/A |
| HQ728265.1 | 197 | IPR038996 | 754 | IPR038993 | N/A | N/A | IPR008258 IPR023346 IPR000189 | N/A | 1332 | IPR038994 |
| KT852574.1 | 201 | IPR038996 | 757 | IPR038993 | N/A | N/A | IPR008258 IPR023346 IPR000189 | N/A | 1326 | N/A |
| OQ507919.1 | 201 | IPR038996 | 759 | IPR038993 | N/A | N/A | IPR008258 IPR023346 IPR000189 | N/A | 1325 | N/A |
| AM183667.1 | 196 | IPR038996 | 760 | IPR038993 | N/A | N/A | IPR008258 IPR023346 IPR000189 | N/A | 1316 | N/A |
| PP079413.1 | 277 | IPR038996 | 817 | IPR038993 | N/A | N/A | IPR013230 | N/A | 1326 | N/A |
| OQ137559.1 | 415 | N/A | 730 | N/A | N/A | N/A | IPR023346 | N/A | 1307 | N/A |
| OR367448.1 | 191 | IPR038996 | 736 | N/A | N/A | N/A | IPR023346 | N/A | 1533 | N/A |
| KT381879.1 | 470 | N/A | 736 | N/A | N/A | N/A | IPR023346 | N/A | 1309 | N/A |
| MK984681.1 | 191 | IPR038996 | 743 | N/A | N/A | N/A | IPR023346 | N/A | 1552 | N/A |

| | | | | | | | | | | |
|---|---|---|---|---|---|---|---|---|---|---|
| ON932081.1 | 288 | N/A | 786 | N/A | N/A | N/A | IPR023346 | N/A | 1368 | N/A |
| PP079414.1 | 289 | N/A | 787 | N/A | N/A | N/A | IPR023346 | N/A | 1367 | N/A |
| MF278336.1 | 179 | IPR038996 | 761 | N/A | N/A | N/A | IPR023346 IPR016047 | N/A | 1957 | N/A |
| MZ592920.1 | 286 | N/A | 742 | N/A | N/A | N/A | IPR023346 IPR023347 IPR002196 | N/A | 1254 | N/A |
| OK042080.1 | 197 | IPR038996 | 750 | IPR038993 | N/A | N/A | IPR023346 IPR023347 IPR002196 IPR000189 | N/A | 1318 | IPR038994 |
| OL856139.1 | 203 | IPR038996 | 752 | IPR038993 | N/A | N/A | IPR023346 IPR023347 IPR002196 IPR000189 | N/A | 1314 | IPR038994 |
| MN988539.1 | 147 | IPR038996 | 955 | N/A | N/A | N/A | IPR052179 IPR003709 | N/A | 1286 | N/A |
| AM265639.1 | 210 | IPR038996 | 855 | N/A | N/A | IPR002196 IPR023347 | N/A | N/A | 1004 | N/A |
| FN594518.1 | 210 | IPR038996 | 861 | N/A | N/A | IPR002196 IPR023347 IPR023346 | N/A | N/A | 1015 | N/A |
| EF160123.1 | 224 | IPR038996 | 952 | N/A | N/A | IPR002196 IPR023347 IPR023346 | N/A | N/A | 1295 | N/A |
| AY288927.2 | 240 | IPR038996 | 979 | N/A | N/A | IPR002196 IPR023347 | N/A | N/A | 1271 | N/A |

| | | | | | | | | | | |
|---|---|---|---|---|---|---|---|---|---|---|
| | | | | | | IPR023346 | | | | |
| KC310802.1 | 315 | N/A | 988 | N/A | N/A | IPR002901 | N/A | N/A | 1403 | |
| HM559717.1 | 414 | N/A | 1035 | N/A | N/A | IPR008258<br>IPR023346<br>IPR016047<br>IPR011055 | N/A | N/A | 1315 | N/A |
| HQ317388.1 | | N/A | 788 | N/A | N/A | IPR023346 | N/A | N/A | | N/A |
| MH540083.1 | 229 | N/A | 826 | N/A | N/A | IPR023346 | N/A | N/A | 1208 | N/A |
| MH729818.1 | 242 | IPR038996 | 981 | N/A | N/A | IPR023346 | N/A | N/A | 1268 | N/A |
| MG845683.1 | 182 | IPR038996 | 887 | N/A | N/A | IPR023346<br>IPR002196 | N/A | N/A | 1311 | N/A |
| MG251390.1 | 241 | IPR038996 | 983 | N/A | N/A | IPR023346<br>IPR002196 | N/A | N/A | 1103 | N/A |
| MK962628.1 | 251 | IPR038996 | 921 | N/A | N/A | IPR023346<br>IPR016047 | N/A | N/A | 1281 | N/A |
| MN844877.1 | 219 | IPR038996 | 928 | N/A | N/A | IPR023346<br>IPR016047 | N/A | N/A | 1187 | N/A |
| KM370384.1 | 190 | IPR038996 | 983 | N/A | N/A | IPR023346<br>IPR016047 | N/A | N/A | 1261 | N/A |
| KM819695.1 | 226 | IPR038996 | 984 | N/A | N/A | IPR023346<br>IPR016047 | N/A | N/A | 1274 | N/A |
| JQ837901.1 | 233 | IPR038996 | 992 | N/A | N/A | IPR023346<br>IPR016047 | N/A | N/A | 1263 | N/A |
| MT104468.1 | 177 | IPR038996 | 874 | N/A | N/A | IPR023346<br>IPR023347<br>IPR002196 | N/A | N/A | 1013 | N/A |

| | | | | | | | | | | |
|---|---|---|---|---|---|---|---|---|---|---|
| MT354570.1 | 177 | IPR038996 | 875 | N/A | N/A | IPR023346<br>IPR023347<br>IPR002196 | N/A | N/A | 1011 | N/A |
| ON148527.1 | 231 | IPR038996 | 979 | N/A | N/A | IPR023346<br>IPR023347<br>IPR002196 | N/A | N/A | 1280 | N/A |
| ON461912.1 | 223 | IPR038996 | 982 | N/A | N/A | IPR023346<br>IPR023347<br>IPR002196 | N/A | N/A | 1267 | N/A |
| MZ398244.1 | 241 | IPR038996 | 982 | N/A | N/A | IPR023346<br>IPR023347<br>IPR002196 | N/A | N/A | 1265 | N/A |
| OX359471.1 | 213 | IPR038996 | 917 | | N/A | IPR023347 | N/A | N/A | 1292 | N/A |
| KC310806.1 | 327 | N/A | 876 | N/A | N/A | IPR041219 | N/A | IPR016047 | 1052 | N/A |
| MK016662.1 | 373 | N/A | 857 | N/A | N/A | IPR042047 | N/A | N/A | 1357 | N/A |
| MT424636.1 | 258 | N/A | 901 | N/A | N/A | IPR042047 | N/A | N/A | 1326 | N/A |
| HQ317389.1 | 239 | N/A | 905 | N/A | N/A | IPR042047 | N/A | N/A | 1322 | N/A |
| OQ999402.1 | 308 | N/A | 998 | N/A | N/A | IPR051056<br>IPR002901 | N/A | N/A | 1405 | N/A |
| MW015080.1 | 286 | N/A | 1053 | N/A | N/A | IPR051056<br>IPR002901<br>IPR011055<br>IPR016047 | N/A | N/A | 1320 | N/A |
| KY117485.1 | 244 | IPR038996 | 916 | N/A | N/A | IPR023346<br>IPR016047 | N/A | N/A | 1274 | N/A |
| AJ505558.1 | 182 | IPR038996 | 899 | N/A | N/A | IPR023346<br>IPR023347 | N/A | N/A | 1338 | N/A |
| OL634959.1 | 204 | IPR038996 | 912 | N/A | N/A | IPR023346 | N/A | N/A | 1001 | N/A |

| | | | | | | | | | | |
|---|---|---|---|---|---|---|---|---|---|---|
| | | | | | | IPR023347 | | | | |
| OQ376857.1 | 189 | IPR038996 | 1061 | N/A | IPR008258 IPR023346 IPR000189 | N/A | N/A | N/A | 1160 | N/A |
| OQ376858.1 | 177 | IPR038996 | 1085 | N/A | IPR008258 IPR023346 IPR000189 | N/A | N/A | N/A | 1131 | N/A |
| JX483874.1 | 173 | IPR038996 | 1080 | N/A | IPR011105 IPR042047 | N/A | N/A | N/A | 1249 | N/A |
| MN988466.1 | 170 | IPR038996 | 1116 | N/A | IPR011105 | N/A | N/A | N/A | 1179 | N/A |
| MT708544.1 | 173 | IPR038996 | 1077 | N/A | IPR042047 IPR011105 | N/A | N/A | N/A | 1293 | N/A |
| MZ462995.1 | 169 | IPR038996 | 1103 | N/A | IPR042047 IPR011105 | N/A | N/A | N/A | 1179 | N/A |
| MT708546.1 | 169 | IPR038996 | 1105 | N/A | IPR042047 IPR011105 | N/A | N/A | N/A | 1182 | N/A |
| MW960034.1 | 174 | IPR038996 | 1120 | N/A | IPR042047 IPR011105 | N/A | N/A | N/A | 1170 | N/A |
| OR997969.1 | 169 | IPR038996 | 1184 | N/A | IPR042047 IPR011105 IPR013230 | N/A | N/A | N/A | 1271 | N/A |
| MF403005.1 | 169 | IPR038996 | 1192 | N/A | IPR042047 IPR011105 | N/A | N/A | N/A | 1255 | N/A |
| OR420753.1 | 187 | IPR038996 | 531 | N/A | N/A | N/A | N/A | N/A | 1206 | N/A |
| OR420755.1 | 179 | IPR038996 | 544 | N/A | N/A | N/A | N/A | N/A | 1346 | N/A |

| | | | | | | | | | | |
|---|---|---|---|---|---|---|---|---|---|---|
| MH598799.1 | 196 | IPR038996 | 558 | N/A | N/A | N/A | N/A | N/A | 1247 | N/A |
| OR464703.1 | 160 | IPR038996 | 567 | N/A | N/A | N/A | N/A | N/A | 1294 | N/A |
| OR420743.1 | 200 | IPR038996 | 568 | N/A | N/A | N/A | N/A | N/A | 1302 | N/A |
| OP329100.1 | 197 | IPR038996 | 571 | N/A | N/A | N/A | N/A | N/A | 1211 | N/A |
| KC465900.1 | 197 | IPR038996 | 609 | N/A | N/A | N/A | N/A | N/A | 1231 | N/A |
| MT375523.1 | 201 | IPR038996 | 615 | N/A | N/A | N/A | N/A | N/A | 1235 | N/A |
| HQ332140.1 | 194 | N/A | 625 | N/A | N/A | N/A | N/A | N/A | 1214 | N/A |
| MZ666938.1 | 206 | IPR038996 | 647 | N/A | N/A | N/A | N/A | N/A | 1269 | N/A |
| KC465901.1 | 168 | IPR038996 | 674 | N/A | N/A | N/A | N/A | N/A | 1266 | N/A |
| ON624112.1 | 200 | IPR038996 | 699 | N/A | N/A | N/A | N/A | N/A | 1297 | N/A |
| MG545917.1 | 204 | IPR038996 | 700 | N/A | N/A | N/A | N/A | N/A | 1249 | N/A |
| KF322026.1 | 202 | IPR038996 | 702 | N/A | N/A | N/A | N/A | N/A | 1230 | N/A |
| MK455769.1 | 204 | IPR038996 | 704 | N/A | N/A | N/A | N/A | N/A | 1260 | N/A |
| KJ502657.1 | 205 | IPR038996 | 706 | N/A | N/A | N/A | N/A | N/A | 1229 | N/A |
| GU071107.1 | 165 | N/A | 708 | N/A | N/A | N/A | N/A | N/A | 1355 | N/A |
| OR420734.1 | 172 | N/A | 708 | N/A | N/A | N/A | N/A | N/A | 1218 | N/A |

| | | | | | | | | | | |
|---|---|---|---|---|---|---|---|---|---|---|
| MW865291.1 | 202 | IPR038996 | 708 | N/A | N/A | N/A | N/A | N/A | 1229 | N/A |
| OM913599.1 | 200 | IPR038996 | 715 | N/A | N/A | N/A | N/A | N/A | 1299 | N/A |
| KJ749827.1 | 202 | IPR038996 | 715 | N/A | N/A | N/A | N/A | N/A | 1288 | N/A |
| MH113813.1 | 192 | IPR038996 | 719 | N/A | N/A | N/A | N/A | N/A | 1258 | N/A |
| MN478376.1 | 221 | IPR038996 | 721 | N/A | N/A | N/A | N/A | N/A | 1289 | N/A |
| MH113812.1 | 194 | IPR038996 | 727 | N/A | N/A | N/A | N/A | N/A | 1259 | N/A |
| KY065149.1 | 180 | N/A | 728 | N/A | N/A | N/A | N/A | N/A | 1684 | N/A |
| MT227925.1 | 204 | IPR038996 | 730 | N/A | N/A | N/A | N/A | N/A | 1413 | N/A |
| MG018929.2 | 189 | IPR038996 | 733 | N/A | N/A | N/A | N/A | N/A | 1275 | N/A |
| EU652770.3 | 190 | IPR038996 | 736 | IPR038993 | N/A | N/A | N/A | N/A | 1313 | N/A |
| HQ641340.1 | 195 | IPR038996 | 738 | IPR038993 | N/A | N/A | N/A | N/A | 1128 | N/A |
| JN991020.1 | 254 | N/A | 739 | N/A | N/A | N/A | N/A | N/A | 1241 | N/A |
| GU071102.1 | 207 | N/A | 742 | N/A | N/A | N/A | N/A | N/A | 1526 | N/A |
| ON755176.1 | 253 | N/A | 743 | N/A | N/A | N/A | N/A | N/A | 864 | N/A |
| KM199771.1 | 182 | IPR038996 | 745 | N/A | N/A | N/A | N/A | N/A | 1242 | N/A |
| MW145136.1 | 236 | IPR038996 | 746 | N/A | N/A | N/A | N/A | N/A | 1456 | N/A |
| MG775261.1 | 251 | IPR038996 | 755 | N/A | N/A | N/A | N/A | N/A | 1278 | N/A |
| KX066068.1 | 251 | IPR038996 | 760 | N/A | N/A | N/A | N/A | N/A | 1194 | N/A |
| AM084414.1 | 196 | IPR038996 | 761 | IPR038993 | N/A | N/A | N/A | N/A | 1296 | N/A |
| HG818824.1 | 197 | IPR038996 | 761 | IPR038993 | N/A | N/A | N/A | N/A | 1299 | N/A |
| KJ183191.1 | 181 | IPR038996 | 768 | N/A | N/A | N/A | N/A | N/A | 1175 | N/A |
| MG878892.2 | 253 | IPR038996 | 774 | N/A | N/A | N/A | N/A | N/A | 1137 | N/A |

| | | | | | | | | | | |
|---|---|---|---|---|---|---|---|---|---|---|
| HG793132.1 | 309 | IPR038996 | 786 | N/A | N/A | N/A | N/A | N/A | 1666 | N/A |
| MT708545.1 | 175 | IPR038996 | 807 | N/A | N/A | N/A | N/A | N/A | 1271 | N/A |
| MW057854.1 | 210 | IPR038996 | 837 | IPR038993 | N/A | N/A | N/A | N/A | 870+468 | N/A |
| EF372997.1 | 235 | N/A | 837 | N/A | N/A | N/A | N/A | N/A | 1416 | N/A |
| AY939843.2 | 201 | N/A | 838 | N/A | N/A | N/A | N/A | N/A | 1246 | N/A |
| HQ316584.1 | 233 | N/A | 852 | N/A | N/A | N/A | N/A | N/A | 1808 | N/A |
| KC310805.1 | 214 | N/A | 858 | N/A | N/A | N/A | N/A | N/A | 1611 | N/A |
| MZ803112.1 | 178 | N/A | 860 | N/A | N/A | N/A | N/A | N/A | 1193 | N/A |
| MK562503.1 | 204 | IPR038996 | 869 | N/A | N/A | N/A | N/A | N/A | 1280 | N/A |
| OM913597.1 | 175 | IPR038996 | 885 | N/A | N/A | N/A | N/A | N/A | 960 | N/A |
| MT259468.2 | 206 | IPR038996 | 887 | N/A | N/A | N/A | N/A | N/A | 992 | N/A |
| MW286266.1 | 201 | IPR038996 | 890 | N/A | N/A | N/A | N/A | N/A | 1250 | N/A |
| MF754111.1 | 250 | IPR038996 | 893 | N/A | N/A | N/A | N/A | N/A | 1285 | N/A |
| GQ413938.2 | 196 | IPR038996 | 895 | N/A | N/A | N/A | N/A | N/A | 1233 | N/A |
| JQ267518.1 | 207 | IPR038996 | 895 | N/A | N/A | N/A | N/A | N/A | 1237 | N/A |
| OQ921331.1 | 189 | IPR038996 | 897 | N/A | N/A | N/A | N/A | N/A | 1257 | N/A |
| KT240186.1 | 203 | IPR038996 | 900 | N/A | N/A | N/A | N/A | N/A | 1262 | N/A |
| MN270887.1 | 204 | IPR038996 | 900 | N/A | N/A | N/A | N/A | N/A | 1231 | N/A |
| JX290549.1 | 205 | IPR038996 | 905 | N/A | N/A | N/A | N/A | N/A | 1264 | N/A |
| KU310944.1 | 181 | IPR038996 | 910 | N/A | N/A | N/A | N/A | N/A | 1265 | N/A |

| | | | | | | | | | | |
|---|---|---|---|---|---|---|---|---|---|---|
| LC776701.1 | 186 | IPR038996 | 910 | N/A | N/A | N/A | N/A | N/A | 1260 | N/A |
| AB854109.1 | 237 | IPR038996 | 910 | N/A | N/A | N/A | N/A | N/A | 1316 | N/A |
| FR687252.1 | 229 | IPR038996 | 915 | N/A | N/A | N/A | N/A | N/A | 1369 | N/A |
| MK387869.1 | 214 | IPR038996 | 918 | N/A | N/A | N/A | N/A | N/A | 1270 | N/A |
| KF319020.1 | 215 | IPR038996 | 918 | N/A | N/A | N/A | N/A | N/A | 1280 | N/A |
| OM513679.2 | 226 | IPR038996 | 922 | N/A | N/A | N/A | N/A | N/A | 1375 | N/A |
| MZ333135.1 | 225 | IPR038996 | 926 | N/A | N/A | N/A | N/A | N/A | 1373 | N/A |
| KF669656.1 | 234 | IPR038996 | 939 | N/A | N/A | N/A | N/A | N/A | 1056 | N/A |
| MN497414.1 | 236 | IPR038996 | 949 | N/A | N/A | N/A | N/A | N/A | 1137 | N/A |
| KR149290.1 | 224 | IPR038996 | 962 | N/A | N/A | N/A | N/A | N/A | 1033 | N/A |
| MT783706.1 | 234 | IPR038996 | 964 | N/A | N/A | N/A | N/A | N/A | 1395 | N/A |
| KJ473423.1 | 235 | IPR038996 | 966 | N/A | N/A | N/A | N/A | N/A | 1050 | N/A |
| HQ337022.1 | 243 | N/A | 967 | N/A | N/A | N/A | N/A | N/A | 1364 | N/A |
| KX397280.1 | 192 | IPR038996 | 1091 | N/A | N/A | N/A | N/A | N/A | 1122 | N/A |
| OQ540924.1 | 170 | N/A | 1301 | N/A | N/A | N/A | N/A | N/A | 1263 | N/A |
| MN857473.1 | 165 | IPR038996 | unclear | N/A | N/A | N/A | N/A | N/A | 1330 | N/A |
| OK570185.1 | 205 | IPR038996 | 712 | N/A | N/A | N/A | N/A | N/A | 1313 | N/A |
| AF338467.2 | 213 | N/A | 1001 | N/A | N/A | N/A | N/A | N/A | 1640 | N/A |
